# A defined multi-kingdom microbial consortium naturalizes the immune system of laboratory mice

**DOI:** 10.64898/2026.10.09.757975

**Authors:** Yi Yang, Shan Liu, Chaoting Zhou, Nile Bayard, Siddharth Krishnamurthy, Elias Roth Gerrick, Michael R. Howitt, Sebastian Shaw, Boris Striepen, Marco Ruella, Ken Cadwell

**Affiliations:** Division of Gastroenterology and Hepatology, Perelman School of Medicine, University of Pennsylvania; Philadelphia, PA 19104, USA; Center for Cellular Immunotherapies, Perelman School of Medicine, University of Pennsylvania; Philadelphia, PA 19104, USA; Division of Hematology and Oncology, Perelman School of Medicine, University of Pennsylvania; Philadelphia, PA 19104, USA; Abramson Cancer Center, University of Pennsylvania; Philadelphia, PA 19104, USA; Cell and Molecular Biology Graduate Program, Perelman School of Medicine, University of Pennsylvania; Philadelphia, PA 19104, USA; The Institute for Immunology and Immune Health, University of Pennsylvania; Philadelphia, PA 19104, USA; Department of Immunology and Microbiology, School of Medicine, University of Colorado Anschutz; Aurora, CO 80045, USA; Department of Pathology, School of Medicine, Stanford University; Stanford, CA 94305, USA; Department of Microbiology and Immunology, School of Medicine, Stanford University; Stanford, CA 94305, USA; Department of Pathobiology, School of Veterinary Medicine, University of Pennsylvania; Philadelphia, PA 19104, USA

## Abstract

Dirty mice experiments suggest that laboratory mice fail to model certain human immune responses due to their limited microbial exposure. The identity, nature, and number of microbes required to normalize the immune system are unknown. Through systematic screening, we uncovered unexpected cross-kingdom microbial interactions and identified four enteric microbes spanning viruses, fungi, and protists sufficient to reproduce key immunological features of pet store mice and adult humans. This consortium also altered cancer outcomes, shifting tumorigenesis toward the anatomical pattern of human familial adenomatous polyposis in *Apc*^Min/+^ mice and recapitulating the imperfect response of many patients to chimeric antigen receptor (CAR) T cell therapy. Thus, surprisingly few microbes can profoundly reshape immune states and disease responses, providing a tractable approach to naturalize laboratory mouse immunity.

## Main Text

Laboratory mice provide powerful models for investigating immune mechanisms and evaluating therapeutic interventions; however, their immune systems differ substantially from those of adult humans. One important contributor is the restricted microbial exposure imposed by specific pathogen-free (SPF) housing, which excludes most infectious agents encountered by free-living mammals. SPF mice consequently display an immunologically immature state characterized by reduced innate immune activation and a predominance of naïve lymphocytes, resembling aspects of human neonatal immunity (*1*, *2*). By contrast, wild and pet store mice exhibit enhanced cytokine signaling, expanded tissue-resident T cells, and environmentally imprinted adaptive immune memory that more closely resemble adult human immunity (*1–3*). Dirty mouse studies have shown that exposing laboratory mice to microbes encountered by wild or pet store mice shift the immune system towards a mature state, promoting innate immune activation and expansion of granulocytes and effector memory-like T cells (*2*, *4–11*). Importantly, such exposure can also enable laboratory mice to reproduce clinically relevant responses to immune-modulating therapies that are poorly modeled under SPF conditions. Mice with enhanced microbial experience recapitulate the proinflammatory response to CD28 superagonist (CD28SA) treatment that caused severe cytokine release syndrome in humans (*11*, *12*) and develop immune checkpoint blockade-associated colitis, a common adverse effect in patients (*13*). Collectively, these findings underscore the importance of microbial exposure in establishing a physiological immune baseline that influences subsequent immune perturbations.

However, the complexity of current dirty mouse models limits their utility. Variation in microbial composition complicates standardization across experiments and laboratories, specialized housing infrastructure is often required, and the contributions of specific microbes to immune and disease phenotypes are difficult to dissect. More fundamentally, these models leave unresolved how much microbial complexity is actually required to produce broad immune maturation. Wild and pet store mice encounter diverse viruses, bacteria, fungi, and other eukaryotic microbes, raising two possibilities: mature immunity may depend on the cumulative effects of extensive microbial exposure, or much of this phenotype may instead be driven by a limited subset of microbes. Resolving this question would clarify the microbial requirements for immune maturation and determine whether the effects of complex microbial experiences can be reproduced in a defined, experimentally tractable system that enables individual microbial contributions to be dissected. We therefore systematically screened a panel of representative infectious microbes, individually and in combination, to determine whether a defined set could recapitulate immune maturation induced by broad microbial exposure and alter disease and therapeutic responses.

### Screening polymicrobial exposures reveals cross-kingdom microbial interactions

The prolonged and laborious experimental time course required for sequential infection represents a major bottleneck for testing multiple microbial combinations in mice. To overcome this limitation, we developed a cohousing-based screening strategy that leverages natural horizontal transmission between mice in the same cage (Fig. 1A). Like previous dirty mouse models (*2*, *6*, *9*, *12*), we used C57BL/6 mice that are commonly used in immunology studies. Adult female mice were individually gavaged with one microbe, and five mice carrying different microbes were then cohoused 7 days later, with each mouse serving as both a source and recipient of microbial transmission. By allowing simultaneous transmission of up to five microbes, this strategy enabled parallel screening of multiple microbial combinations rather than testing one defined cocktail at a time.

**Fig. 1.**
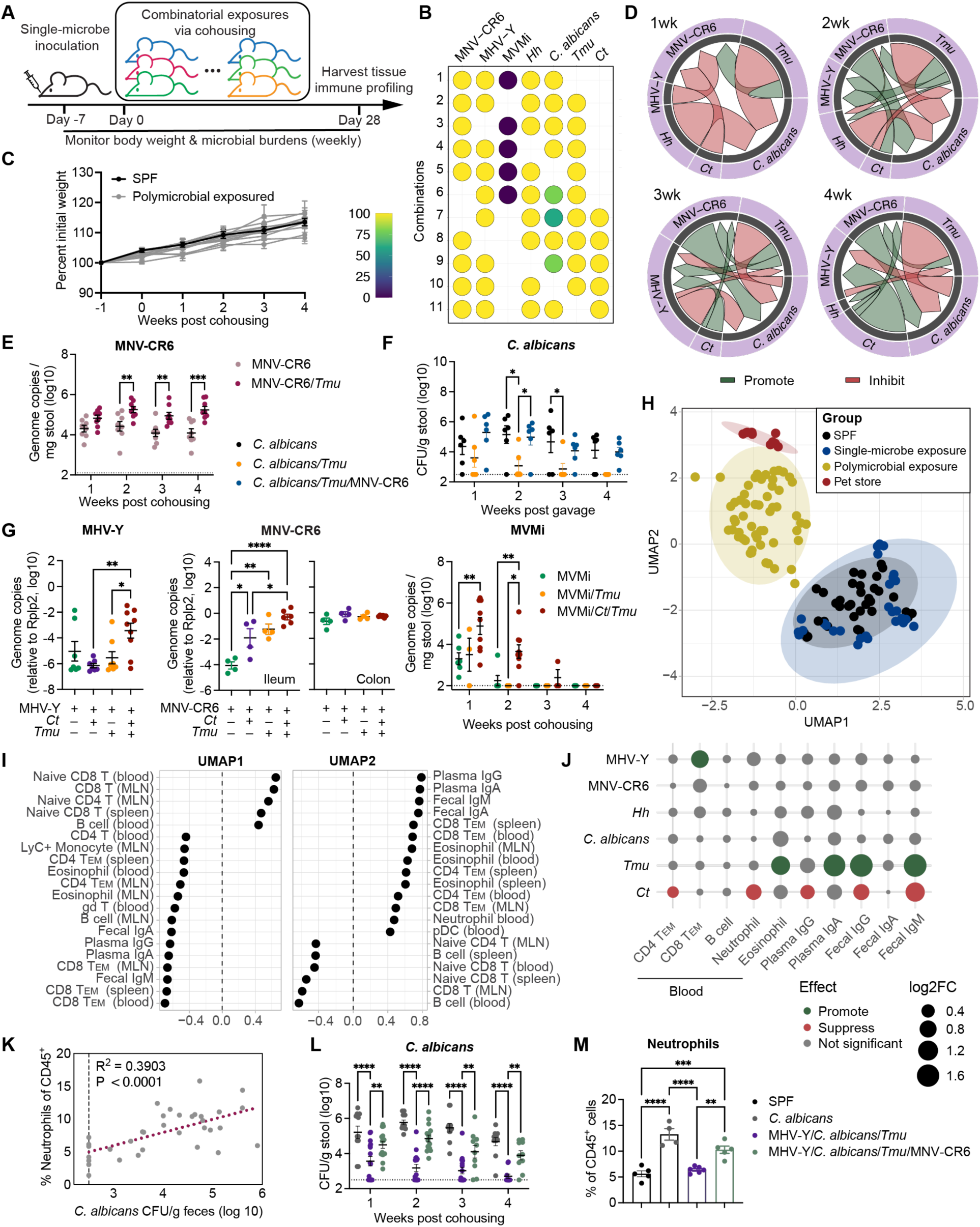
Systematic microbial screening identifies a defined four-microbe consortium that drives immune maturation. (**A**) Experimental design for combinatorial microbial screening. (**B**) Microbial transmission frequencies during cohousing, determined from fecal microbial burdens. (**C**) Body weight after microbial exposure. Each line represents one microbial combination. (**D**) Directed pairwise microbial interactions based on microbial burdens in combinations with or without the indicated second microbe. Arrow width indicates log10 fold change. Only interactions with *P* < 0.05 by unpaired t test are shown. (**E**) MNV-CR6 fecal burden in MNV-CR6-infected and MNV-CR6/*Tmu*-coinfected mice. n = 8 mice per group. (**F**) *C. albicans* fecal burden following inoculation in SPF mice or mice preinfected with *Tmu* or *Tmu*/MNV-CR6. n = 6 mice per group. (**G**) MHV-Y burden in colonic epithelial fractions (2 weeks), MNV-CR6 burden in ileal and colonic epithelial fractions (7 weeks), and MVMi fecal burden following indicated microbial exposures. n = 3 to 9 mice per group. (**H**) UMAP of immune profiles across microbial exposure conditions. Immune features (see fig. S3) were assessed at 4 weeks post cohousing. (**I**) Top 20 immune features correlated with UMAP1 and UMAP2 coordinates by Spearman correlation. (**J**) Associations between individual microbes and immune features. Mice carrying each microbe were compared with mice lacking that microbe across all tested combinations. Dot size represents absolute log2 fold change (log2FC) of immune features between microbe-positive and microbe-negative mice. (**K**) Correlation between *C. albicans* fecal burden and blood neutrophils among *C. albicans*-exposed mice without *Ct* at 4 weeks post cohousing. Dots represent individual mice. (**L** and **M**) *C. albicans* fecal burden (n = 11 to 16 mice per group) (L) and blood neutrophils (n = 4 to 6 mice per group) (M) following indicated microbial exposures. Data are pooled from two independent experiments (E to G, and L to M) and are presented as mean ± SEM (C, E to G, and L to M). Dotted lines in (E to G), and (L) denote limit of detection. Statistical analysis was performed using two-way analysis of variance (ANOVA) with Sidak correction (E), or one-way (F, G, and M) or two-way (L) ANOVA with Tukey post hoc test. \**P* < 0.05; \*\**P* < 0.01; \*\*\**P* < 0.001; \*\*\*\**P* < 0.0001.

We selected candidate microbes that are prevalent in wild or pet store mice but depleted from SPF mouse colonies. Although previous dirty mouse models involve microbial exposures with diverse tissue tropisms, enteric microbes constitute the majority of transmissible agents detected in these models (*2*, *7*, *14*, *15*), leading us to hypothesize that intestinal exposures alone may be sufficient to drive immune maturation. We therefore focused on enteric candidates and selected seven microbes spanning viruses, bacteria, fungi, and protists with distinct immunological effects that collectively cover major branches of immunity, including Th1, Th2, and Th17 responses (*16–21*) (fig. S1A). All these agents were originally isolated from rodents except the fungal component. Because SPF mice are neutropenic relative to adult humans (*22*), and our previous studies showed that fungal exposure contributes to neutrophil expansion in rewilded mice (*6*) and that enteric *Candida albicans* colonization promotes granulopoiesis in laboratory mice (*23*), we included the CHN1 strain, which efficiently colonizes SPF mice without antibiotic pretreatment (*24*). Seven microbes yielded 21 possible five-microbe combinations, providing a feasible screening space while retaining substantial microbial diversity.

We first monitored fecal microbial burdens longitudinally following gavage with individual microbes (fig. S1B). All seven microbes successfully infected or colonized SPF mice and reached peak burden within 1 to 2 weeks. Murine norovirus CR6 strain (MNV-CR6), *Helicobacter hepaticus* (*Hh*), *C. albicans*, *Cryptosporidium tyzzeri* (*Ct*, STL strain), and *Tritrichomonas musculus* (*Tmu*) remained detectable at 4 weeks, whereas minute virus of mice i strain (MVMi) largely declined beginning at 2 weeks and murine hepatitis virus Y strain (MHV-Y) became undetectable in most mice by 4 weeks. No overt signs of disease were observed (fig. S1C), suggesting that these microbes represent pathobionts and commensals under these conditions.

We next generated polymicrobial exposures through cohousing. To minimize variation in baseline microbiota and facilitate reproducibility across studies, all mice in the screen were obtained from The Jackson Laboratory. Body weight and microbial burdens were monitored weekly, and blood, spleen, and mesenteric lymph nodes (MLN) were collected 4 weeks after cohousing (Fig. 1A). MNV-CR6, MHV-Y, *Hh*, *Tmu*, and *Ct* transmitted efficiently, whereas *C. albicans* transmission varied across combinations, suggesting context-dependent colonization and spread (Fig. 1B and fig. S2A). MVMi was detected only in initially inoculated mice and was therefore excluded from subsequent experiments. Polymicrobial exposures caused no overt disease or loss of body weight (Fig. 1C and fig. S2B).

In total, we evaluated 11 five-microbe combinations (table S1) across 55 mice, generating 275 longitudinal fecal samples and 110 pairwise microbial co-occurrence contexts. Network analysis identified multiple interactions among the tested microbes (Fig. 1D). In one example, *Tmu* was associated with increased MNV-CR6 burden across microbial combinations (fig. S2C), consistent with reports that *Tmu* induces intestinal tuft cell hyperplasia (*21*, *25*) and that tuft cells serve as a reservoir for persistent MNV-CR6 infection (*26*). This effect was confirmed by direct comparison of MNV-CR6 infection with or without *Tmu* (Fig. 1E).

We further examined two unexpected three-way interactions identified in polymicrobial exposure settings. First, across the microbial combinations tested, *C. albicans* levels were reduced in the presence of *Tmu* (fig. S2D), consistent with its reduced transmission when the two microbes coexisted in the same cage (Fig. 1B and fig. S2A). The combinatorial screen further revealed that the presence of MNV-CR6 restored *C. albicans* levels in mice exposed to *Tmu* (fig. S2D). We validated this interaction by directly testing the three implicated microbes (Fig. 1F). Second, coinfection with the protists *Ct* and *Tmu* enhanced all three viruses in our screen. MHV-Y and MNV-CR6 burdens were higher in mice exposed to both protists than in mice without this combination, and although *Tmu* alone increased MNV-CR6 burden, the addition of *Ct* further amplified this effect (fig. S2E). An independent validation experiment confirmed that *Ct* and *Tmu* together increased MHV-Y and MNV-CR6 levels beyond those observed with either protist alone, with the effect on MNV-CR6 particularly pronounced in the ileum (Fig. 1G). Moreover, MVMi, which otherwise failed to transmit during cohousing, successfully transmitted in the presence of *Ct* and *Tmu*, reached higher fecal burdens, and exhibited delayed clearance (Fig. 1G). These results reveal the potential for higher-order, cross-kingdom interactions to occur during polymicrobial exposures.

### Polymicrobial exposure drives systemic immune maturation

We next asked whether exposures to combinations of gut microbes broadly reshape the immune system by quantifying major immune cell populations in blood, spleen, and MLN, together with immunoglobulin levels in plasma and feces 4 weeks after cohousing (fig. S3). The analysis included SPF mice and mice exposed to individual microbes or polymicrobial combinations, with adult pet store mice serving as a reference. Pet store mice were purchased from a local pet shop and confirmed to be outbred *Mus musculus* by genetic monitoring. To visualize relationships among immune profiles across microbial exposure conditions, we applied Uniform Manifold Approximation and Projection (UMAP) to 65 immune readouts from 152 mice (Fig. 1H). SPF mice and single-microbe-exposed mice clustered closely, indicating limited immune remodeling in response to individual microbes. However, polymicrobial exposures induced a marked shift toward the immune profile of pet store mice, suggesting that broad systemic immune maturation could be achieved with exposure to a small set of microbes.

### Complementary immune activity and microbial compatibility guide consortium construction

The substantial variation in immune profiles across different polymicrobial exposures suggested that the specific composition of microbial exposure strongly influences the resulting immune state and could potentially be optimized to maximize immune maturation. We therefore examined which immune features contributed most strongly to the major axes of separation in the UMAP analysis. Correlating individual immune variables with UMAP coordinates identified key features associated with UMAP1 and UMAP2, including CD62L^+^CD44^−^ naïve and CD62L^−^CD44^+^ effector memory (T_EM_) CD8 T cells, CD4 T_EM_ cells, eosinophils, neutrophils, and B cells across tissues, and immunoglobulin levels in plasma and feces (Fig. 1I). Notably, many of these features are also altered in dirty mouse models generated through complex microbial exposure (*2*, *5*, *11*).

We hypothesized that effective immune maturation may require microbes with complementary immunological activities that collectively induce distinct features of the mature immune state. Therefore, we examined associations between individual microbes and the immune features contributing to UMAP separation. MHV-Y was associated with expansion of CD8 T_EM_ cells, with higher CD8 T_EM_ frequencies observed across combinations containing MHV-Y (Fig. 1J and fig. S4A). *Tmu* was associated with increased eosinophils and elevated immunoglobulin levels (Fig. 1J and fig. S4B), consistent with a recent study identifying *Tmu* as a natural adjuvant that promotes mucosal IgA production (*27*). In contrast, *Ct* infection was negatively associated with several immune features, including CD4 T_EM_ cells, neutrophils, plasma and fecal IgG, and fecal IgM (Fig. 1J and fig. S4C). The previously described ability of *C. albicans* to promote neutrophil expansion (*23*) was not apparent in the initial analysis (Fig. 1J). Because *Ct* was negatively associated with neutrophil abundance, we reasoned that the presence of *Ct* might obscure the relationship between *C. albicans* and neutrophil expansion. Indeed, after excluding *Ct*-infected mice, blood neutrophil frequency positively correlated with fecal *C. albicans* burden (Fig. 1K). These findings indicate that the immunomodulatory effects associated with individual microbes depend on the broader microbial context. Together, MHV-Y, *Tmu*, and *C. albicans* emerged as complementary phenotype-driving microbes associated with distinct features of immune maturation.

Based on these complementary immune activities, we tested a three-microbe combination comprising MHV-Y, *C. albicans*, and *Tmu*. Polymicrobial exposure was established using the same cohousing strategy as in the initial screen. MHV-Y and *Tmu* followed their expected infection kinetics; however, *C. albicans* failed to colonize effectively, and the expected neutrophil expansion was not observed (Fig. 1, L and M). This result was consistent with our earlier finding that *Tmu* restricts *C. albicans* colonization (Fig. 1F and fig. S2D), demonstrating that complementary immunological activity alone was insufficient to ensure an effective consortium.

The earlier polymicrobial screen also revealed that MNV-CR6 counteracted *Tmu*-mediated restriction of *C. albicans* colonization (Fig. 1F). We therefore hypothesized that adding MNV-CR6 to the three-microbe combination would restore *C. albicans* colonization. Indeed, within the resulting four-microbe consortium, *C. albicans* colonization was restored and was accompanied by neutrophil expansion (Fig. 1, L and M). Thus, MNV-CR6 enabled the immune activity elicited by *C. albicans* by resolving an incompatibility between microbial components.

We next characterized the four-microbe consortium comprising MHV-Y, MNV-CR6, *C. albicans*, and *Tmu*. Following cohousing, all four microbes successfully infected or colonized SPF mice (fig. S5A). MNV-CR6 and *Tmu* persisted at high levels in feces for up to 12 weeks, whereas MHV-Y and *C. albicans* reached robust levels initially but became undetectable in feces beginning approximately 4 weeks after cohousing. Immune profiling at 4 weeks showed that mice exposed to this four-microbe consortium were clearly separated from SPF mice and clustered closer to pet store mice (fig. S5B). As with the other polymicrobial combinations tested, exposure to this four-microbe consortium caused no body weight loss or overt signs of disease (fig. S5C). Notably, mice exposed to the four-microbe consortium alone clustered closer to pet store mice than mice exposed to combinations containing these same four microbes plus *Hh*, MVMi, or *Ct* (fig. S5B). Thus, increasing the number of microbial exposures did not necessarily enhance immune maturation, and the four-microbe combination was sufficient to largely reproduce the broad immune remodeling observed with more complex microbial exposures.

### IMPACT4 recapitulates the systemic immune features of pet store mice

We defined this four-microbe consortium as IMPACT4 (Immune-Modulatory PAthobionts and Commensals, Tetra-mix) and next characterized the breadth and kinetics of the immune remodeling it induced. Mice exposed to this consortium (hereafter, IMPACT4 mice) are generated using the cohousing strategy. Longitudinal profiling of circulating immune populations following IMPACT4 exposure revealed rapid but dynamic changes across both innate and adaptive compartments (Fig. 2A). During the first week, multiple myeloid populations expanded, including natural killer (NK) cells, granulocytes, plasmacytoid dendritic cells (pDCs), conventional dendritic cells (cDCs), and Ly6C^+^ monocytes, accompanied by a transient decrease in the relative frequency of lymphocytes. Most myeloid populations contracted after the second week, whereas neutrophils and eosinophils remained elevated for 4 to 8 weeks. B cell proportions remained reduced, while CD4 and CD8 T cells recovered and expanded by 4 weeks post cohousing.

**Fig. 2.**
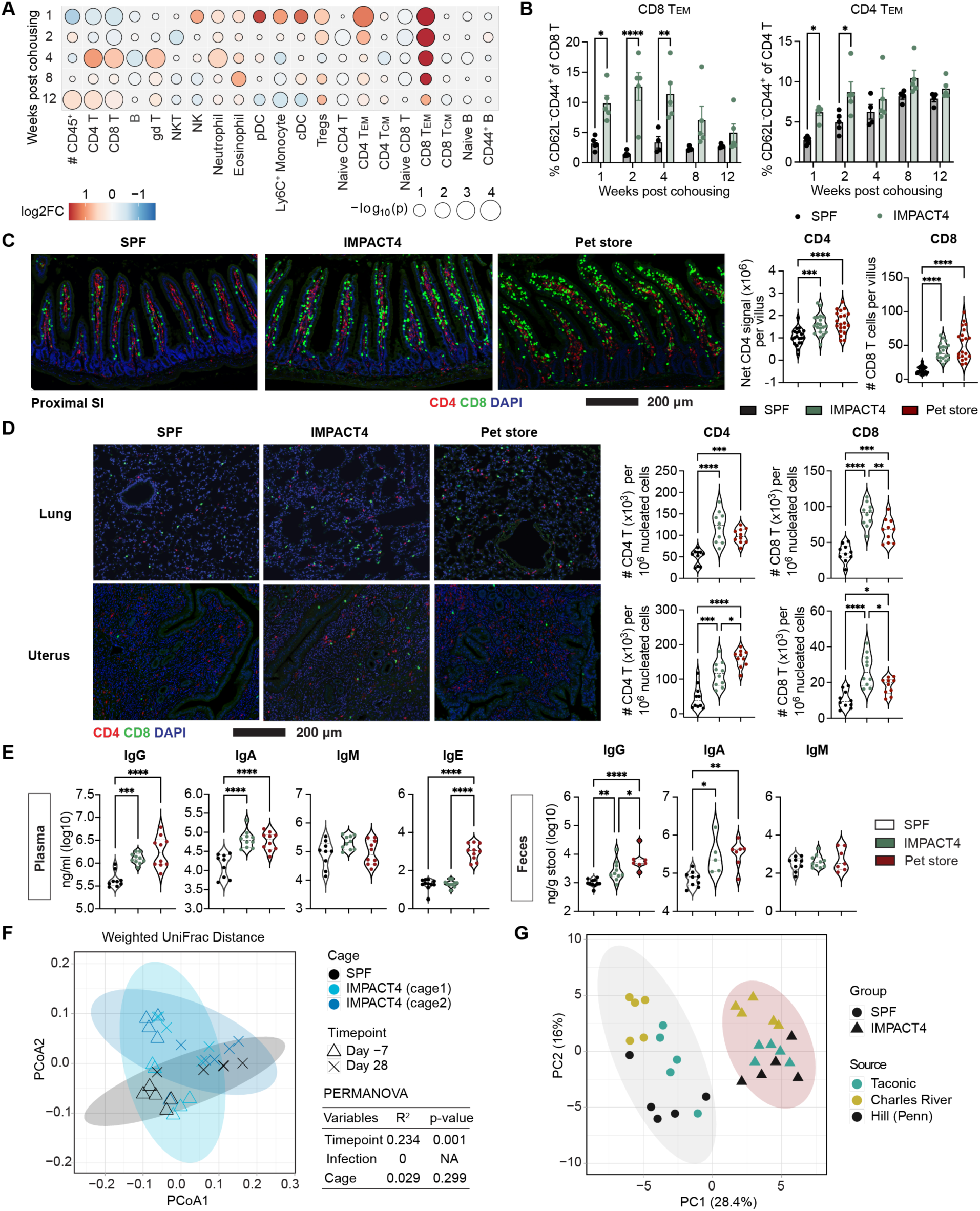
IMPACT4 recapitulates systemic immune maturation observed in pet store mice. (**A**) Longitudinal changes in blood immune populations following IMPACT4 exposure. Log2 fold change (log2FC) relative to SPF controls was calculated from group mean values. (**B**) Blood CD8 and CD4 T_EM_ cells in SPF (n = 4 to 5) and IMPACT4 (n = 5) mice. Data are presented as mean ± SEM. (**C** and **D**) Immunofluorescence staining and quantification of CD4 and CD8 T cells in proximal small intestine (SI) (C), lung, and uterus (D) from SPF, IMPACT4, and pet store mice. T cell density was quantified from 20 intestinal villi or 10 randomly selected fields using at least two mice per group. Representative images are shown. (**E**) Plasma and fecal immunoglobulin levels in SPF (n = 9), IMPACT4 (n = 10), and pet store (n = 10) mice. (**F**) Beta diversity of fecal microbiota from SPF (n = 5) and two cages of IMPACT4 mice (n = 5 mice per cage) before and after microbial exposure. Contributions of experimental variables were assessed by PERMANOVA. (**G**) Principal component analysis (PCA) of immune profiles in SPF and IMPACT4 mice from different sources. Data are pooled from (E) or are representative of two independent experiments (A to D). Violin plots (C to E) indicate median and quartiles. Statistical analysis was performed using unpaired t test (A), two-way ANOVA with Sidak correction (B), or one-way ANOVA with Tukey post hoc test (C to E). \**P* < 0.05; \*\**P* < 0.01; \*\*\**P* < 0.001; \*\*\*\**P* < 0.0001.

Within the T cell compartment, IMPACT4 induced a prominent and sustained expansion of effector memory-like populations (Fig. 2, A and B). CD8 T_EM_ cells, a hallmark of previously described dirty mouse models (*2*, *6*, *11*), increased as early as the first week after cohousing and remained elevated through 8 weeks, with the increase persisting at 12 weeks in a subset of mice. CD4 T_EM_ cells also increased during the first 2 weeks and remained elevated thereafter, although less prominently than CD8 T_EM_ cells (Fig. 2B). More detailed phenotyping at 4 weeks showed that changes in circulating T cells were largely confined to the antigen-experienced CD44^+^ compartment, consistent with previous comparisons between pet store and laboratory mice (*2*) (fig. S6A). CD8 T cells from IMPACT4 mice showed increased KLRG1 and reduced CD62L, CD127, CD27, and CXCR3, consistent with enhanced effector differentiation. CD4 T cells similarly showed reduced CD62L and CD27 within the CD44^+^ population. In contrast, cell size (FSC), proliferation (Ki67), and markers of recent antigen exposure (PD-1 and CD69) were unchanged, indicating that IMPACT4 establishes a more differentiated antigen-experienced T cell pool at this timepoint rather than driving sustained T cell activation or proliferation.

We next asked whether T cell remodeling extended beyond the circulation. Immunofluorescence staining revealed increased CD8 T cell density throughout the intestinal tract of IMPACT4 mice, resembling the distribution observed in pet store mice (Fig. 2C and fig. S6B). CD4 T cell density was also increased in both IMPACT4 and pet store mice, but this increase was largely restricted to the proximal small intestine (Fig. 2C and fig. S6B). Although exclusively consisting of enteric microbes, IMPACT4 increased CD4 and CD8 T cell densities beyond the intestine, including in the lung, uterus, skin, liver, and kidney (Fig. 2D and fig. S6C). Thus, a defined enteric microbial exposure was sufficient to remodel tissue-localized T cell compartments both locally and systemically.

These cellular changes were accompanied by enhanced humoral responses, as reflected by increased total IgG and IgA in both plasma and feces compared with SPF mice, reaching levels comparable to those in pet store mice (Fig. 2E). IgM levels were similar across groups, whereas elevated IgE was observed only in pet store mice, potentially reflecting differences in genetic background or parasitic exposures not represented by IMPACT4. Together, these data indicate that exposure to IMPACT4 drives coordinated remodeling of innate, T cell, and humoral compartments, establishing a systemic immune state that closely resembles that of pet store mice.

### IMPACT4-induced immune maturation is robust across microbiota contexts

The endogenous microbiota plays an important role in shaping host immunity (*28*). Thus, we asked whether IMPACT4 alters the resident microbiota and whether its immunological effects depend on baseline microbiota composition. We performed 16S rRNA sequencing on fecal samples collected before (day −7) and 4 weeks after IMPACT4 exposure (day 28). Unexposed SPF controls and two independent cages of IMPACT4 mice were included to distinguish temporal and cage-dependent variation from changes associated with IMPACT4 exposure.

Overall changes in microbiota composition following IMPACT4 exposure were modest, as measured by alpha and beta diversity (Fig. 2F and fig. S7A). Permutational multivariate analysis of variance (PERMANOVA) identified timepoint, rather than IMPACT4 exposure or cage, as the major variable contributing to microbiota variation (Fig. 2F). Temporal changes included reductions in Clostridiales and Peptostreptococcales-Tissierellales and expansion of Lachnospirales (fig. S7B). Only two taxa, *Bifidobacterium* sp. and *Turicibacter* sp., were specifically associated with IMPACT4 exposure (fig. S7, C and D). Similar reductions in these taxa have been reported following *Tmu* colonization and attributed to competition for dietary microbiota-accessible carbohydrates (*16*). These results suggest that IMPACT4 exposure did not broadly remodel the resident microbiota, with the limited changes observed consistent with known effects of *Tmu* colonization.

We next examined whether IMPACT4 could induce similar immune remodeling across mice with distinct baseline microbiota. C57BL/6 mice obtained from Taconic Biosciences and Charles River, as well as a C57BL/6 line bred onsite at the University of Pennsylvania, all exhibited shifts in immune profiles away from their corresponding unexposed SPF controls along principal component 1 (PC1) following IMPACT4 exposure (Fig. 2G). Moreover, along PC2, IMPACT4 mice from different sources clustered more closely together than their unexposed counterparts. These findings demonstrate that IMPACT4 induces consistent immune remodeling across distinct mouse sources and that exposure may also reduce baseline immune differences associated with mouse origin.

### Single-step oral gavage extends the utility of IMPACT4

Although cohousing provided an efficient strategy for screening microbial combinations, simultaneous administration of the complete consortium offers a more practical approach for routine use. Concurrent gavage of all four microbes was only partially successful when mice were housed individually (fig. S8A). In contrast, when multiple inoculated mice were housed together, all animals became productively infected with the complete consortium, and each microbe exhibited longitudinal patterns of fecal microbial burdens comparable to those observed following the cohousing approach (fig. S8, A and B). Thus, direct inoculation combined with subsequent cage-level transmission provided a reliable one-step method for establishing IMPACT4 exposure.

Our initial cohousing experiments were restricted to female mice because adult males separated after weaning cannot reliably be cohoused due to aggressive behavior. The one-step gavage approach eliminated the need to introduce unfamiliar animals into the same cage, enabling IMPACT4 exposure in both sexes. Male and female mice showed comparable immune maturation following IMPACT4 exposure (fig. S8C), further extending the applicability of the system.

### IMPACT4 enhances the translational fidelity of laboratory mice

To assess whether IMPACT4 exposure promotes an immune state more akin to that of adult humans, we performed bulk RNA sequencing of whole blood from SPF and IMPACT4 mice and compared their transcriptional profiles. Gene set enrichment analysis (GSEA) revealed broad enrichment of immune-related pathways in IMPACT4 mice, including leukocyte migration, innate and inflammatory responses, cytokine signaling, and humoral immunity (fig. S9), aligning with the systemic immune remodeling observed above. We next applied the analytical framework established in previous studies (*2*) to compare IMPACT4-induced transcriptional changes with signatures of human immune maturation. Using GSEA, we compared genes differentially expressed between IMPACT4 and SPF mice against ranked transcriptomic signatures derived from peripheral blood mononuclear cells (PBMC) of adult and neonatal humans. Genes enriched in IMPACT4 mice showed significant overlap with adult human PBMC signatures, whereas genes enriched in SPF mice were more strongly associated with neonatal human signatures (Fig. 3A). These results indicate that IMPACT4 exposure shifts the immune transcriptional landscape of laboratory mice toward a more adult human-like state.

**Fig. 3.**
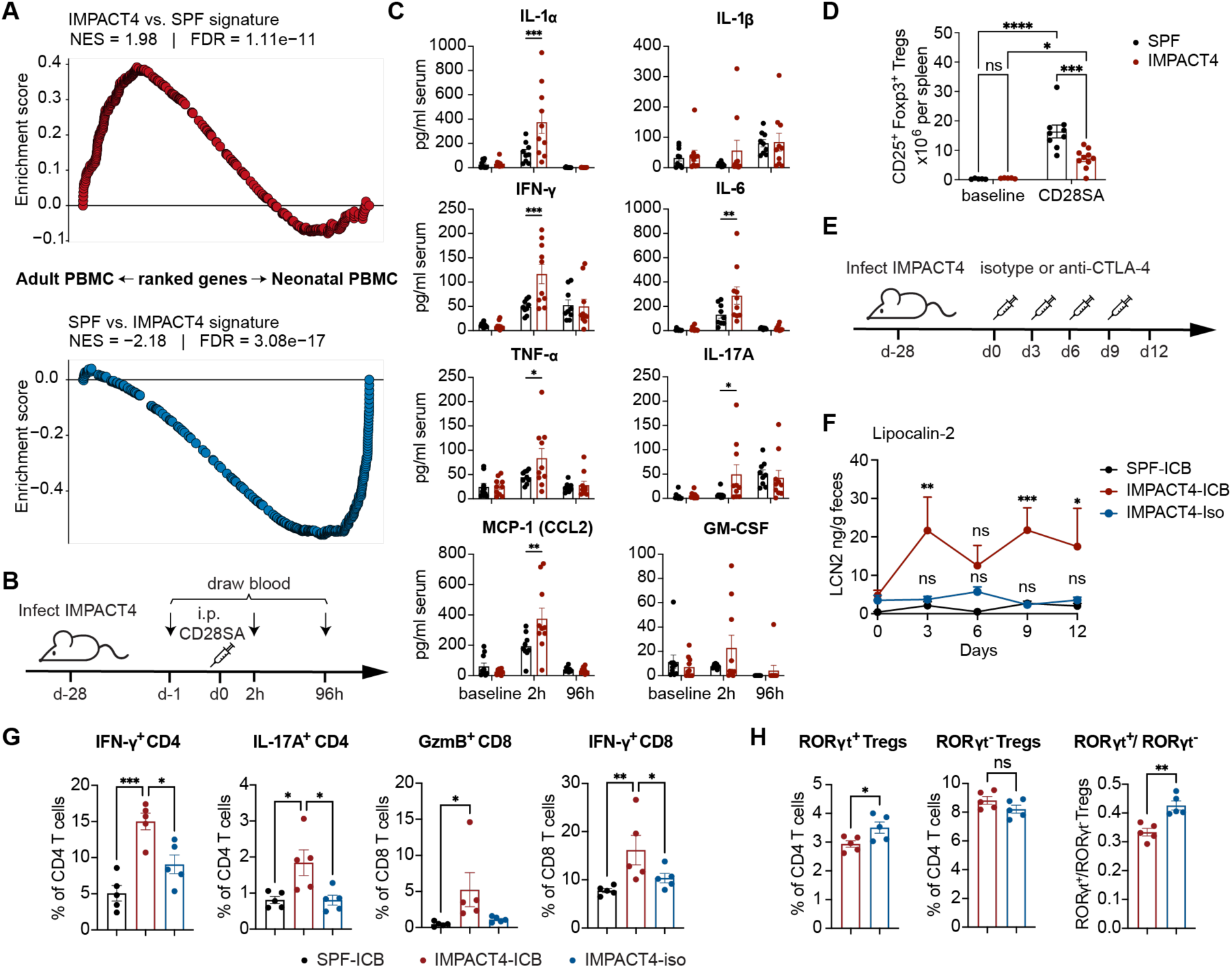
IMPACT4 enhances adult human-like immune features and clinically relevant treatment responses. (**A**) GSEA comparing blood transcriptional signatures from SPF and IMPACT4 mice with human adult versus cord blood PBMC signatures. Mouse signatures comprised the top 400 differentially expressed genes upregulated in each group and mapped to human orthologs. (**B** to **D**) Responses to CD28SA treatment in SPF and IMPACT4 mice. Data are pooled from two independent experiments. (B) Experimental design. (C) Serum cytokine levels at baseline and 2 and 96 hours after CD28SA treatment. n = 10 mice per group. (D) Splenic CD25^+^Foxp3^+^ Tregs at baseline (n = 5 mice per group) and 96 hours (n = 10 mice per group) after CD28SA treatment. (**E** to **H**) Responses to anti-CTLA-4 treatment in SPF and IMPACT4 mice. Data are representative of two independent experiments. (E) Experimental design. (F) Fecal lipocalin-2 (LCN2) levels. n = 5 mice per group. (**G** and **H**) Flow cytometric analysis of cecal lymphocytes 12 days after initiation of anti-CTLA-4 or isotype treatment. n = 5 mice per group. GzmB, granzyme B. Data are presented as mean ± SEM (C, D, and F to H). Statistical analysis was performed using two-way ANOVA with Sidak correction (C and D) or Dunnett correction (versus IMPACT4-Iso) (F), one-way ANOVA with Tukey post hoc test (G), or unpaired t test (H). \**P* < 0.05; \*\**P* < 0.01; \*\*\**P* < 0.001; \*\*\*\**P* < 0.0001; ns, not significant.

Limited microbial exposure in SPF mice contributes to the poor translation of preclinical findings to humans (*29*), and dirty mouse models more faithfully reproduce adult human immune responses across multiple experimental settings (*7*, *12*, *13*, *15*, *30*, *31*). To determine whether IMPACT4 similarly improves translational fidelity, we first examined the response to CD28SA, which induces robust regulatory T cell (Treg) expansion in SPF mice but caused severe cytokine release syndrome in humans (*32*, *33*). Following the experimental framework established in “wildling mice” (*12*), SPF and IMPACT4 mice received a single dose of CD28SA antibody, and serum cytokines were measured at baseline and 2 and 96 hours after treatment (Fig. 3B). Cytokine levels were largely comparable between groups at baseline. At 2 hours after treatment, however, IMPACT4 mice exhibited markedly increased concentrations of multiple proinflammatory cytokines, including interleukin-1α (IL-1α), IL-6, interferon-γ (IFN-γ), tumor necrosis factor-α (TNF-α), IL-17A, and monocyte chemoattractant protein-1 (MCP-1) (Fig. 3C). This cytokine surge was transient and largely resolved by 96 hours. Consistent with this altered response, splenic Treg expansion following CD28SA treatment was substantially reduced in IMPACT4 mice compared with SPF controls (Fig. 3D).

We further tested whether IMPACT4 mice recapitulate immune checkpoint blockade-associated colitis, a common immune-related adverse event in patients (*34*) that is poorly reproduced in SPF mice. Using the experimental framework established in mice reconstituted with wild microbiota (WildR) (*13*), we found that anti-CTLA-4 treatment increased fecal lipocalin-2 in IMPACT4 mice but not in similarly treated SPF mice or IMPACT4 mice receiving isotype control antibody (Fig. 3, E and F). Immune profiling of cecal tissues further revealed that anti-CTLA-4 treatment in IMPACT4 mice was associated with increased frequencies of IFN-γ⁺ and IL-17A⁺ effector T cell populations and reduced anti-inflammatory RORγt⁺ Tregs (Fig. 3, G and H), consistent with observations in WildR mice (*13*). These findings collectively support that IMPACT4 exposure enables laboratory mice to more faithfully recapitulate immune features and clinically relevant treatment-associated toxicities observed in adult humans.

### IMPACT4 shifts *Apc*^Min/+^ tumorigenesis toward the anatomical pattern of human FAP

We then examined whether IMPACT4 exposure could alter outcomes in disease models with well-established differences from human disease. We first focused on intestinal tumorigenesis in *Apc*^Min/+^ mice, a widely used model of familial adenomatous polyposis (FAP) in which *Apc* loss drives constitutive Wnt/β-catenin signaling and spontaneous intestinal adenoma formation (*35–37*). Unlike human FAP, however, tumors in *Apc*^Min/+^ mice arise predominantly in the small intestine rather than the colon, and extensive small intestinal tumor burden leads to progressive morbidity and premature mortality (*35*, *38*). Because host–microbe interactions strongly influence intestinal tumorigenesis, we asked whether IMPACT4 exposure could alter this disease trajectory.

*Apc*^Min/+^ mice were exposed to IMPACT4 beginning at 6 weeks of age or maintained under SPF conditions. IMPACT4 exposure significantly prolonged survival in both sexes, with a greater effect in females (Fig. 4A and fig. S10A). At 16 weeks, IMPACT4 mice exhibited markedly fewer ileal polyps than SPF littermates in both females and males (Fig. 4, B and C, and fig. S10B). Reduced tumor burden was already evident at 10 weeks of age, indicating that IMPACT4 alters tumor development before advanced disease (Fig. 4D and fig. S10C). We then examined Wnt/β-catenin signaling, the major oncogenic pathway driving tumorigenesis in *Apc*^Min/+^ mice. Expression of the Wnt target gene *Axin2* was lower in both tumor-free and tumor-rich ileal regions of IMPACT4 mice (Fig. 4E). Immunohistochemistry similarly revealed reduced β-catenin staining in morphologically normal epithelium and adenomas (Fig. 4F), together with reduced epithelial proliferation as measured by Ki-67 staining (Fig. 4G).

**Fig. 4.**
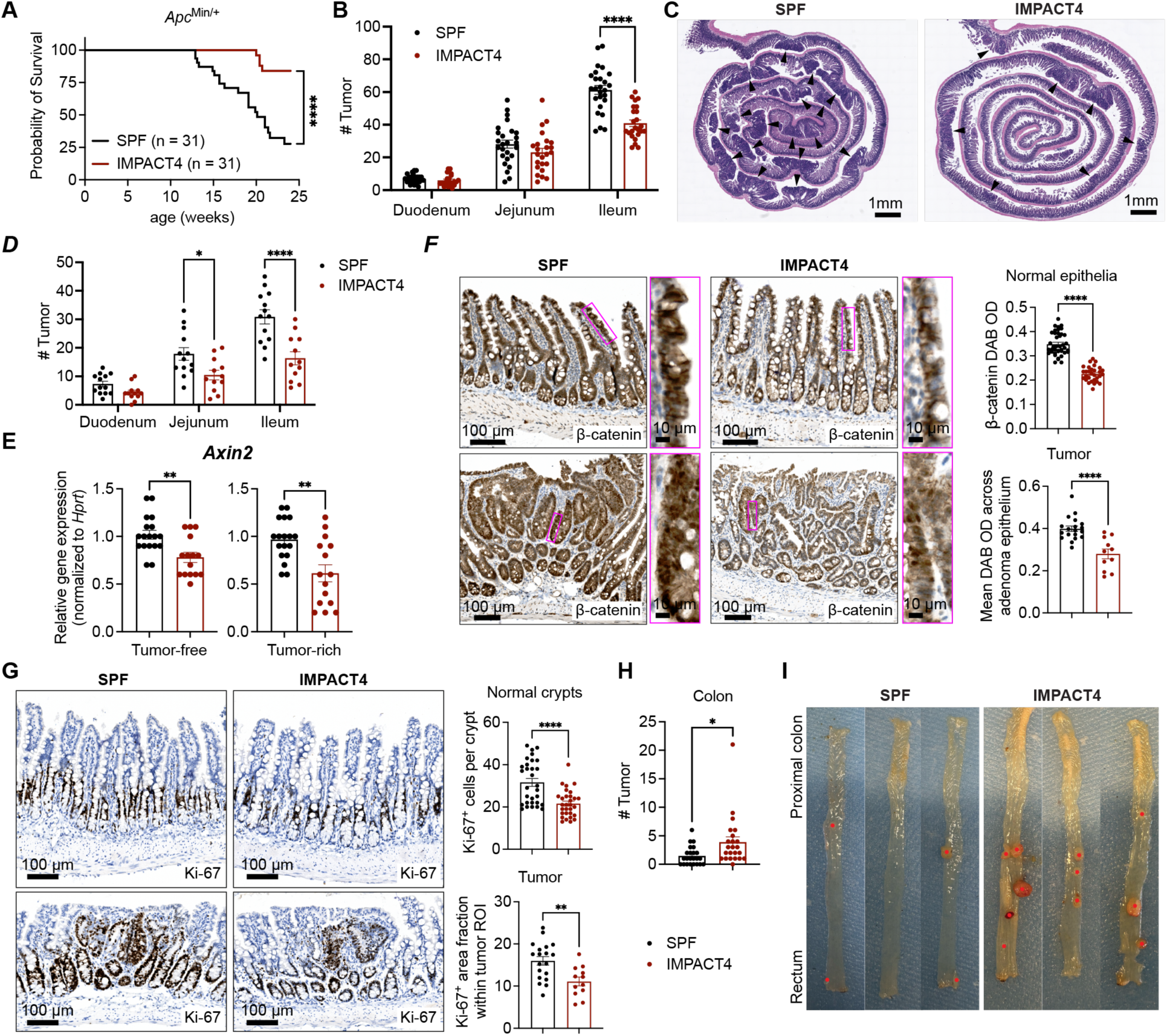
IMPACT4 alters the anatomical distribution of tumors in *Apc*^Min/+^ mice. (**A**) Survival of *Apc*^Min/+^ mice maintained under SPF conditions or exposed to IMPACT4 at 6 weeks of age. n = 31 mice per group. (**B**) Polyp numbers in intestinal segments of 16-week-old *Apc*^Min/+^ mice. n = 26 SPF and 24 IMPACT4 mice. (**C**) Representative H&E-stained ileal Swiss rolls from 16-week-old *Apc*^Min/+^ mice. Arrowheads indicate polyps. (**D**) Polyp numbers in intestinal segments of 10-week-old *Apc*^Min/+^ mice. n = 13 SPF and 12 IMPACT4 mice. (**E**) *Axin2* expression in tumor-free and tumor-rich ileal regions from 10-week-old *Apc*^Min/+^ mice measured by qPCR. Two to three regions of each type were dissected per mouse. n = 18 tissue samples from 7 SPF mice and 15 tissue samples from 6 IMPACT4 mice. (**F** and **G**) Representative β-catenin (F) and Ki-67 (G) immunohistochemical staining and quantification of morphologically normal epithelium and tumors in ileal tissues from 10-week-old *Apc*^Min/+^ mice. Normal epithelium was quantified by β-catenin DAB optical density or Ki-67^+^ cells per crypt using 10 randomly selected villus-crypt units or crypts per mouse (n = 4 SPF and 3 IMPACT4 mice). Tumors were quantified by β-catenin DAB optical density or Ki-67^+^ area fraction. Visible polyps in each group were analyzed (n = 19 SPF and 12 IMPACT4 polyps). DAB, 3,3′-diaminobenzidine; OD, optical density; ROI, region of interest. (**H**) Colonic tumor numbers in male *Apc*^Min/+^ mice at 10 to 20 weeks of age. n = 23 mice per group. (**I**) Representative images of colons in 16-week-old male *Apc*^Min/+^ mice. Red dots indicate tumors. Data are pooled from at least two independent experiments. Data are presented as mean ± SEM (B, D to H). Statistical analysis was performed using log-rank Mantel-Cox test (A), two-way ANOVA with Sidak correction (B and D), or unpaired t test (E to H). \**P* < 0.05; \*\**P* < 0.01; \*\*\**P* < 0.001; \*\*\*\**P* < 0.0001.

In contrast to the suppression of small intestinal tumorigenesis, IMPACT4 exposure increased both the number and size of colonic tumors in male *Apc*^Min/+^ mice (Fig. 4, H and I). This increase was not observed in female mice (fig. S10D). Thus, IMPACT4 exposure shifted the anatomical distribution of intestinal tumorigenesis in male *Apc*^Min/+^ mice toward that observed in human FAP, while also prolonging survival and suppressing small intestinal tumor development in both sexes.

### IMPACT4 limits CART19 therapeutic responses

CAR T cell therapy has achieved major clinical success, but therapeutic responses in patients remain heterogeneous, with many patients showing limited or transient responses (*39*). By contrast, CAR T cells often exhibit robust antitumor activity in SPF mouse models (*40*). We therefore asked whether IMPACT4 exposure could reproduce this clinically relevant limitation of CAR T cell therapy in a syngeneic B cell lymphoma model.

To distinguish adoptively transferred from endogenous T cells, CD45.2^+^ BALB/c recipient mice were maintained under SPF conditions or exposed to IMPACT4, and CD45.1^+^ mice were used to generate CD19-directed CAR T (CART19) cells. Recipients were implanted with CD19^+^ A20 lymphoma cells and treated with CART19 or untransduced (UTD) control T cells following cyclophosphamide preconditioning (Fig. 5A). Tumor growth before treatment was comparable between SPF and IMPACT4 mice (fig. S11A). CART19 treatment markedly suppressed tumor growth under SPF conditions but failed to control tumors in IMPACT4 mice, which grew similarly to those in UTD-treated controls (Fig. 5, B and C, and fig. S11B). Consistent with impaired tumor control, CART19 treatment improved survival in SPF but not IMPACT4 mice (Fig. 5D).

**Fig. 5.**
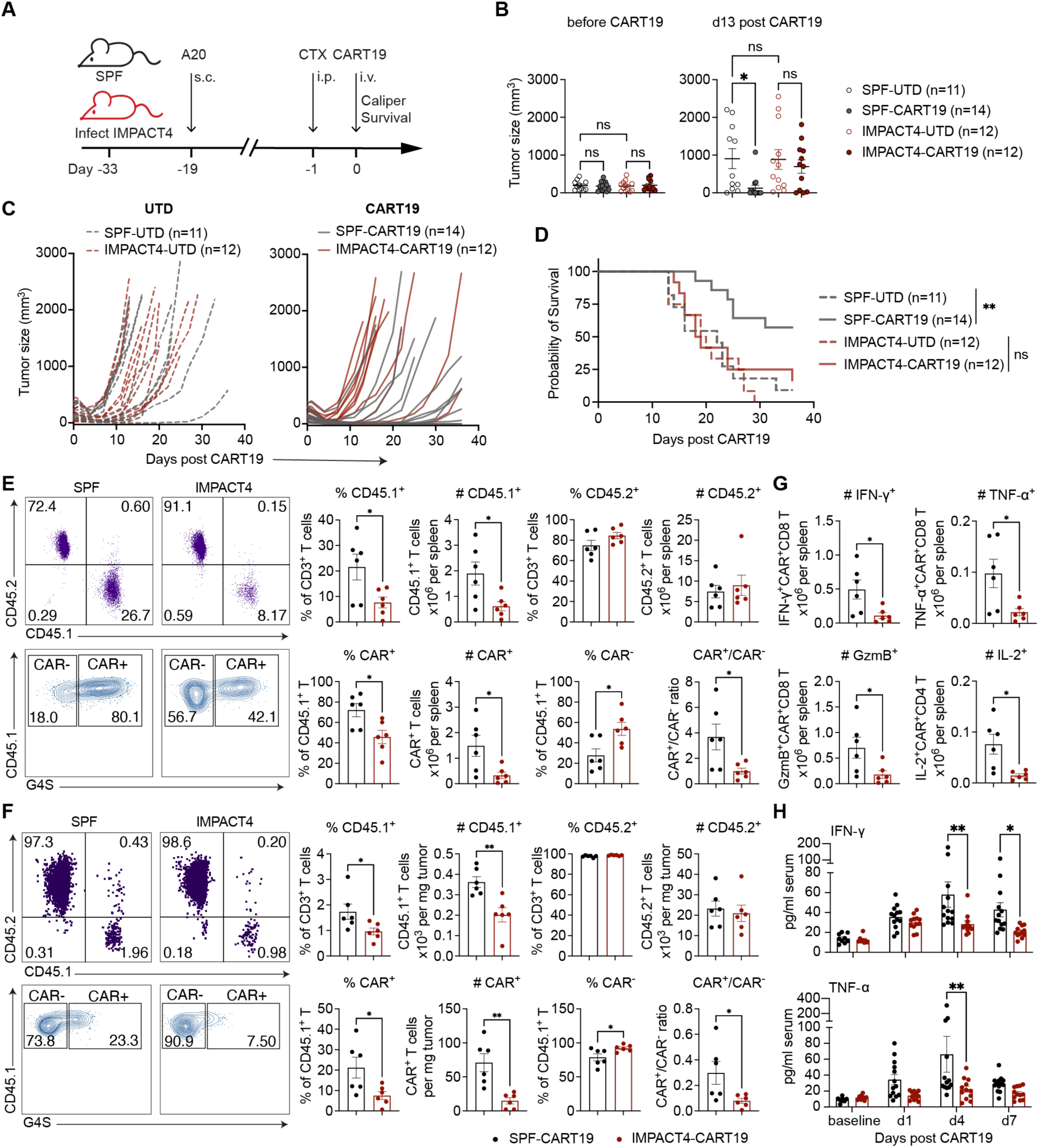
IMPACT4 limits CART19 therapeutic responses. (**A**) Experimental design. CTX, cyclophosphamide. (**B** to **D**) Tumor growth and survival following UTD or CART19 transfer. n = 11 SPF and 12 IMPACT4 mice for UTD treatment; n = 14 SPF and 12 IMPACT4 mice for CART19 treatment. (B) Tumor volumes 1 day before and 13 days after cell transfer. (C) Individual tumor growth curves. (D) Overall survival. (**E** and **F**) Immune profiling of CAR T cells in spleen (E) and tumor (F) 6 days after CART19 transfer. n = 6 mice per group. Representative flow cytometry plots are shown. Cells were pre-gated on CD3^+^ T cells (top) or CD45.1^+^ T cells (bottom). (**G**) Absolute numbers of IFN-γ-, TNF-α-, and granzyme B (GzmB)-producing CAR^+^ CD8 T cells and IL-2-producing CAR^+^ CD4 T cells in spleen 6 days after CART19 transfer. n = 6 mice per group. (**H**) Serum IFN-γ and TNF-α at baseline (n = 9 SPF and 8 IMPACT4 mice) and 1, 4, and 7 days (n = 13 SPF and 12 IMPACT4 mice) after CART19 treatment. Data are pooled from two independent experiments (B to D, and H) and are presented as mean ± SEM (B, and E to H). Statistical analysis was performed using one-way ANOVA with Tukey post hoc test (B), log-rank Mantel-Cox test (D), unpaired t test (E to G), or two-way ANOVA with Sidak correction (H). \**P* < 0.05; \*\**P* < 0.01; ns, not significant.

To investigate the cellular basis of this loss of efficacy, we analyzed donor T cells in the spleen and tumor 6 days after infusion. IMPACT4 mice had fewer CD45.1^+^ donor T cells in the spleen, whereas endogenous CD45.2^+^ T cell abundance was largely unchanged (Fig. 5E). Because the infused product contained both CAR^+^ and CAR^−^ donor T cells (*41*), we further compared these populations within the same host environment. CAR^+^ cells were disproportionately reduced in IMPACT4 mice, resulting in a lower CAR^+^/CAR^−^ donor T cell ratio (Fig. 5E). A similar pattern was also observed among tumor-infiltrating lymphocytes (TILs). IMPACT4 mice had fewer CD45.1^+^ and CAR^+^ T cells in the tumor, together with a lower CAR^+^/CAR^−^ donor T cell ratio (Fig. 5F). IMPACT4 mice consequently had fewer effector CAR^+^ T cells, including IFN-γ^+^, TNF-α^+^, and granzyme B^+^ CD8 T cells, as well as IL-2^+^ CD4 T cells (Fig. 5G). Consistent with these cellular changes, the increase in serum IFN-γ and TNF-α following CART19 infusion was markedly blunted in IMPACT4 mice (Fig. 5H). Collectively, these findings identify microbial exposure as a host determinant of CAR T cell efficacy and associate impaired tumor control with reduced CAR T cell accumulation in both systemic and tumor compartments and attenuated effector responses.

## Discussion

Studies of wild, pet store, and other dirty mice demonstrated that microbial exposures profoundly reshape the immune system of laboratory mice, but the microbial components responsible for these effects have remained poorly defined. Here, we show that broad immune maturation can be reconstructed with a defined combination of only four enteric microbes spanning viruses, fungi, and protists. Exposure to the IMPACT4 consortium remodeled innate and adaptive immune compartments across multiple tissues, reproduced key features associated with adult human immunity, and better reflected human responses to clinically relevant treatments. These findings suggest that the immune consequences of complex microbial experience can emerge from a limited set of functionally distinct microbial exposures.

The screening process further revealed that the effects of polymicrobial exposure depend strongly on the identity and interactions of the microbes involved. Three individual IMPACT4 members were associated with distinct immunological effects, including expansion of CD8 T_EM_ with MHV-Y infection, increased immunoglobulin production and eosinophil expansion with *Tmu*, and neutrophil expansion in the presence of *C. albicans*. Rather than contributing directly to these immune features, MNV-CR6 counteracted *Tmu*-mediated restriction of *C. albicans*, thereby facilitating its colonization and sustaining *C. albicans*-associated immune changes. These complementary activities provide a potential explanation for how a small consortium can generate broad immune remodeling. Moreover, the additional microbes tested in this study did not further enhance immune maturation beyond that achieved by the four-member IMPACT4 consortium. Adding *Hh* and MVMi provided little further enhancement, whereas including *Ct* even shifted the overall immune profile away from that of pet store mice. Thus, greater microbial diversity alone does not necessarily produce a more physiologically mature immune state.

Notably, all four IMPACT4 members are enteric microbes, suggesting that intestinal microbial exposures are sufficient to drive systemic immune remodeling across multiple tissues. This finding is consistent with the established role of the intestinal mucosa as a major site for immune education, and local microbial encounters can influence distal immunity through immune cell trafficking, soluble mediators, microbially derived metabolites, and microbial translocation (*28*, *42*, *43*). Microbial exposures at other barrier sites, including the respiratory and urogenital tracts, may similarly promote systemic immune maturation but pose greater risks of host morbidity when exposing these sites to multiple microbes. For example, cohousing with pet store mice often causes morbidity and mortality during the initial exposure period (*2*), and sequential infection with pathogens such as influenza requires prolonged periods for recovery (*44*). By contrast, IMPACT4 induced substantial immune maturation without detectable adverse health consequences at baseline. The ability to establish a mature immune baseline while preserving overall host health may be particularly valuable for studies in which preexisting morbidity or tissue injury could confound subsequent disease or therapeutic responses.

The functional consequences of IMPACT4 exposure were evident across multiple disease and therapeutic settings. IMPACT4 reproduced responses previously observed in other dirty mouse models, including heightened proinflammatory responses to CD28 superagonism and immune checkpoint blockade. Importantly, IMPACT4 also shifted outcomes in two cancer models toward clinically relevant features that are poorly captured under SPF conditions. *Apc*^Min/+^ mice predominantly develop small intestinal tumors, in contrast to the colorectal tumor distribution characteristic of human FAP. IMPACT4 suppressed this characteristic small intestinal tumorigenesis in both sexes while increasing colonic tumor development in males, shifting the anatomical pattern of tumorigenesis toward that of human FAP. Similarly, CART19 therapy produced robust tumor control in SPF mice, whereas IMPACT4 exposure reduced therapeutic efficacy and recapitulated the limited response observed in many patients. Together, these findings suggest that incorporating defined microbial experience can reveal clinically relevant disease and therapeutic responses that are obscured in immunologically immature SPF mice.

The microbes examined here represent only a subset of possible physiological exposures, and other combinations may generate similar or distinct mature immune states. The unexpected sufficiency of four microbes nevertheless provides an opportunity to reduce a highly complex environmental variable to experimentally manipulable components. Individual consortium members can be removed, replaced, or modified, allowing the microbial and immunological determinants of mature immune states to be dissected within a reproducible exposure framework. IMPACT4 therefore complements models based on pet store mice, wild mice, and other complex microbial exposures by bridging the biological relevance of broad microbial experience with the experimental control of conventional laboratory models.

## Acknowledgments

We thank Dan R. Littman for generously sharing the bacterial stock of *Helicobacter hepaticus*; former Cadwell lab member Simone Dallari for generating the MVMi stocks; all members of the Cadwell lab for discussions and comments; the University of Pennsylvania Gnotobiotic and Naturalized Mouse Core (Penn-CHOP Microbiome Program, RRID:SCR_022384) for assistance with germ-free mice; Center for Molecular Studies in Digestive and Liver Diseases (P30DK050306) and the Molecular Pathology and Imaging Core (RRID:SCR_022420) for tissue embedding, slicing and H&E staining; Penn Vet Comparative Pathology Core (RRID:SCR_022438) at the University of Pennsylvania for their technical assistance with the IF and IHC staining; Stuart N. Isaacs, and University Laboratory Animal Resources (ULAR) for assistance with animal husbandry design and compliance; and CHOP Flow Cytometry Core for their assistance and resources. Yi Yang, PhD was supported by a Cancer Research Institute Irvington Postdoctoral Fellowship (CRI5231).

## Author contributions

Conceptualization: YY, KC

Methodology: YY, KC, SL, MR, SK, ERG, MRH, SS, BS

Investigation: YY, SL, CZ, NB

Visualization: YY

Funding acquisition: KC

Project administration: YY, KC

Supervision: KC

Writing – original draft: YY, KC

Writing – review & editing: YY, KC, SL, CZ, NB, ERG, MRH, SS, BS

## Data, code, and materials availability

Sequencing data are available at NCBI BioProject: PRJNA1494533. Bulk RNA sequencing (GEO: GSE338940) raw and processed data are available at Gene Expression Omnibus (GEO) and associated raw FASTQ files at Sequence Read Archive (SRA). 16S rRNA sequencing raw FASTQ files are available at SRA. This study did not generate original code. Any additional information required to reanalyze the data reported in this paper is available from the lead contact upon request.

---

**Fig. S1.**
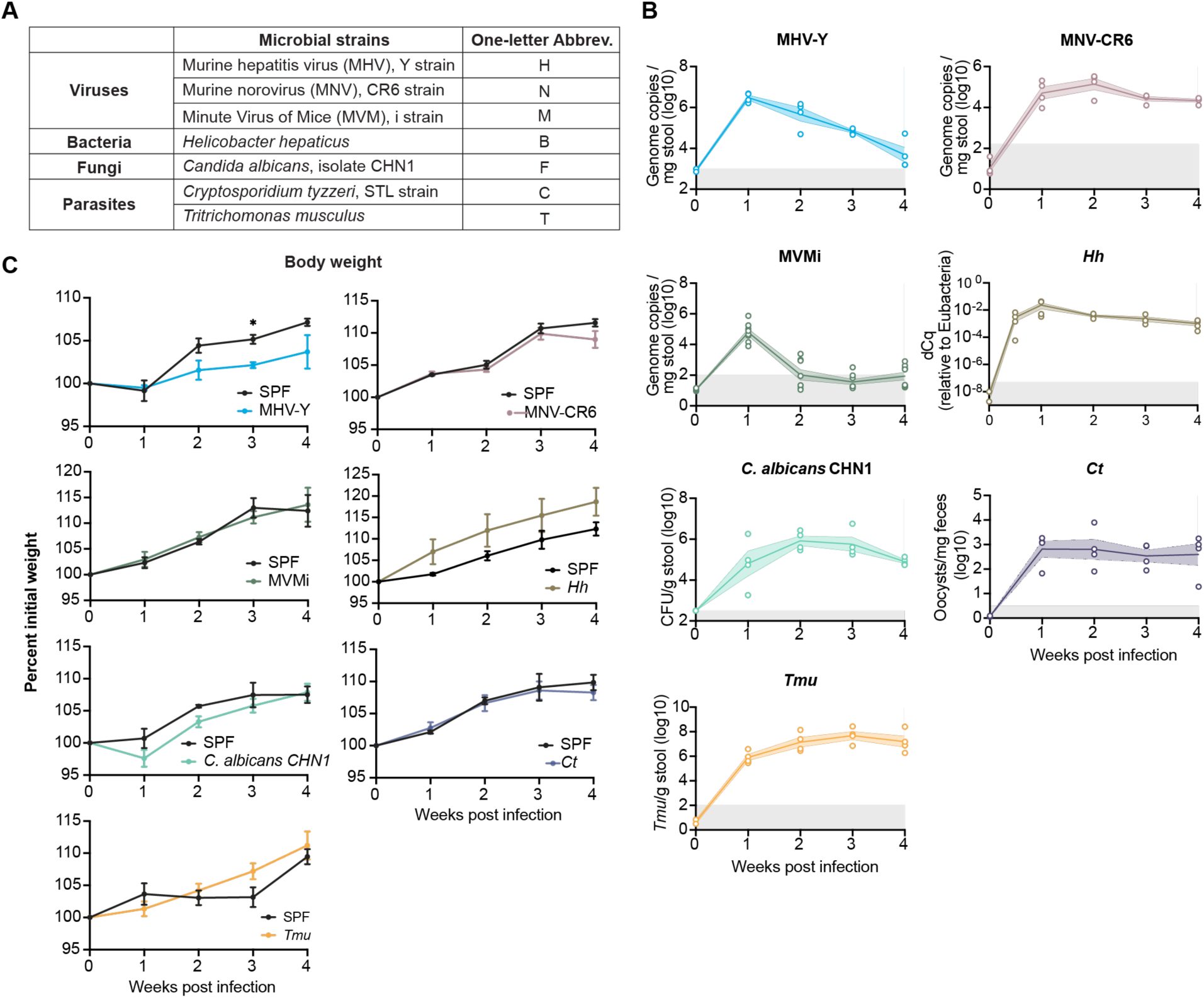
Characterization of individual microbial exposures. (**A**) Taxonomy of the seven microbes selected for screening. (**B**) Longitudinal microbial burdens following single-microbe exposure. Fecal microbial burdens were quantified weekly after gavage. n = 4 to 5 mice per group. Gray shading indicates the limit of detection. (**C**) Body weight of single-microbe-exposed mice and experiment-matched SPF controls. Data are presented as mean ± SEM (B and C). Statistical analysis was performed using two-way repeated-measures ANOVA with Sidak correction at individual time points when a significant interaction was detected (C). \**P* < 0.05.

**Fig. S2.**
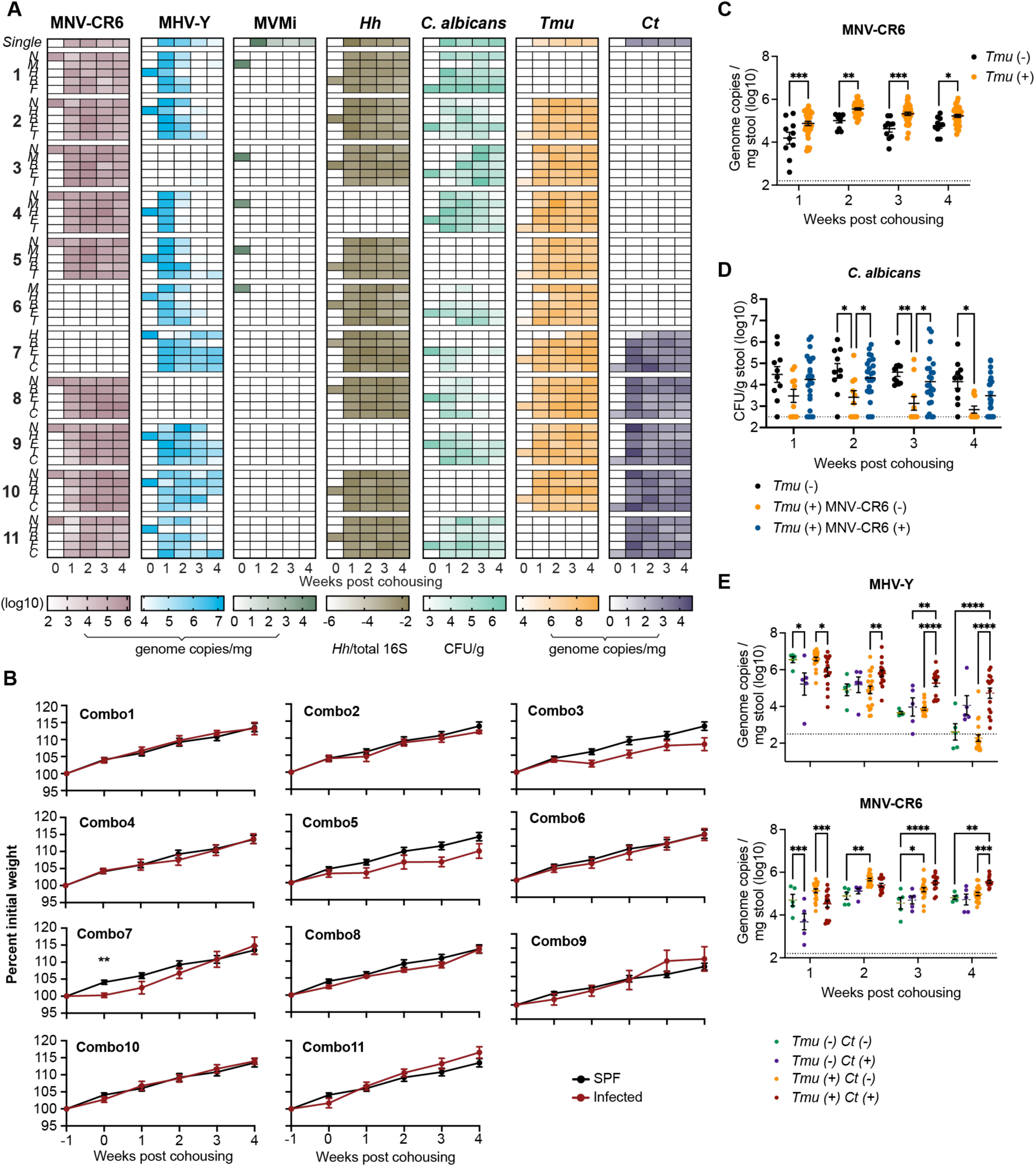
Screening of polymicrobial exposures through cohousing. (**A**) Heatmaps showing longitudinal microbial burdens across tested combinations. Columns represent timepoints and rows represent individual mice, grouped by microbial combination. Numbers indicate the corresponding microbial combination, and letters indicate the microbe used for initial infection of each mouse. Microbial abbreviations are defined in fig. S1A. The top row of each heatmap shows microbial burden during single-microbe infection. (**B**) Body weight of SPF mice and mice exposed to each microbial combination. n = 5 mice per group. (**C**) MNV-CR6 fecal burden in mice exposed to microbial combinations with (n = 35 mice) or without (n = 10 mice) *Tmu*. (**D**) *C. albicans* fecal burden in mice exposed to microbial combinations without *Tmu* (n = 10 mice), with *Tmu* but not MNV-CR6 (n = 10 mice), or with *Tmu* and MNV-CR6 (n = 25 mice). (**E**) Fecal burdens of MHV-Y (top) and MNV-CR6 (bottom) in mice exposed to microbial combinations lacking both *Tmu* and *Ct* (n = 5 mice), containing *Ct* but not *Tmu* (n = 5 mice), containing *Tmu* but not *Ct* (n = 20 mice), or containing both *Tmu* and *Ct* (n = 15 mice). Data are presented as mean ± SEM (B to E). Dotted lines indicate the limit of detection (C to E). Statistical analysis was performed using two-way repeated-measures ANOVA with Sidak correction at individual time points when a significant interaction was detected (B), two-way ANOVA with Sidak correction (C) or Tukey post hoc test (D and E). \**P* < 0.05; \*\**P* < 0.01; \*\*\**P* < 0.001; \*\*\*\**P* < 0.0001.

**Fig. S3.**
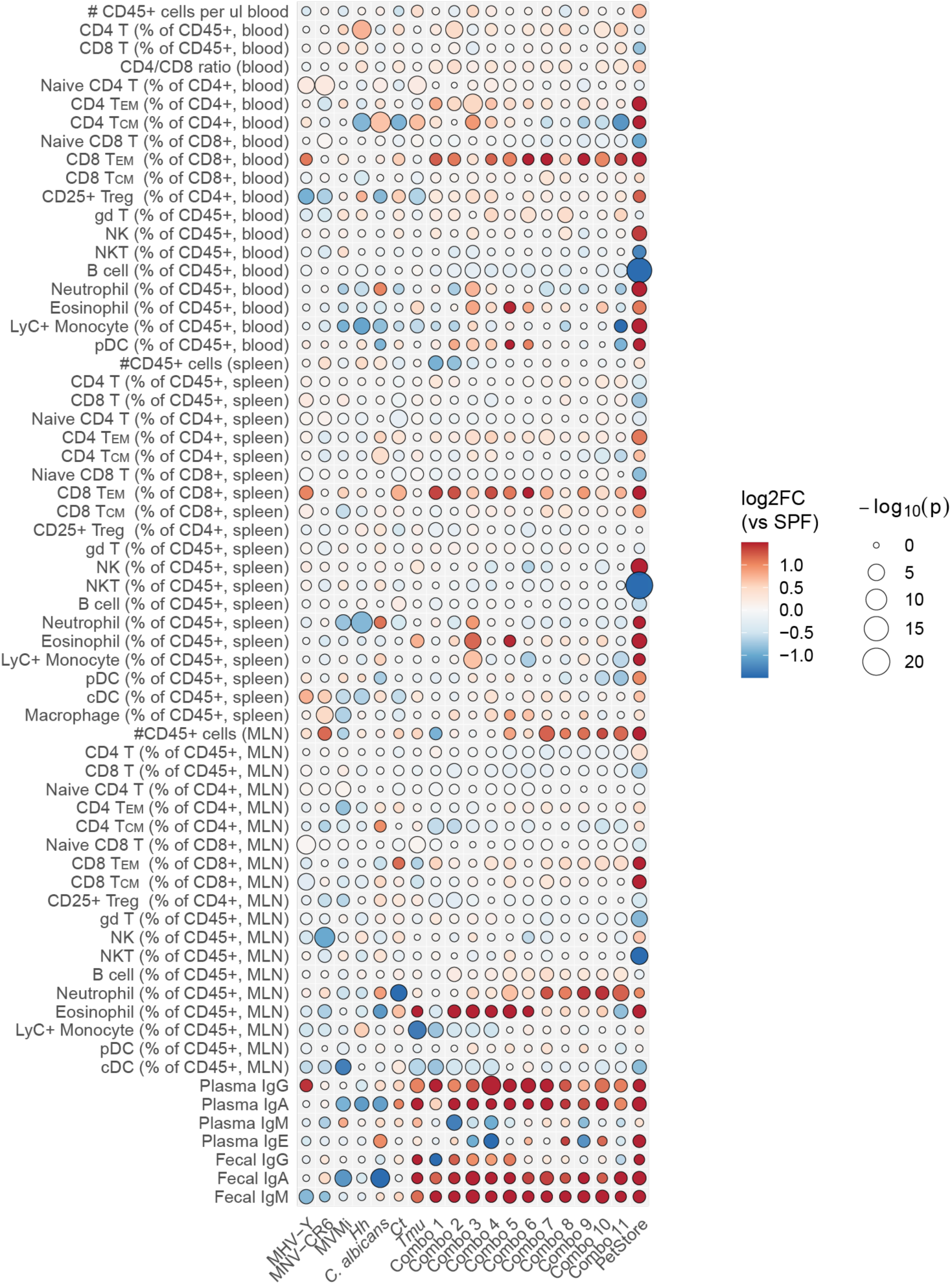
Immune remodeling across individual and polymicrobial exposures. Dot heatmap summarizing changes in immune readouts across mice exposed to individual microbes or defined polymicrobial combinations relative to SPF controls. Columns represent microbial exposure conditions and rows represent immune readouts. Dot color indicates the direction and magnitude of change, calculated as log2 fold change (log2FC) relative to SPF controls from group mean values. Dot size indicates statistical significance as −log10(P value). Statistical significance was determined by unpaired t test.

**Fig. S4.**
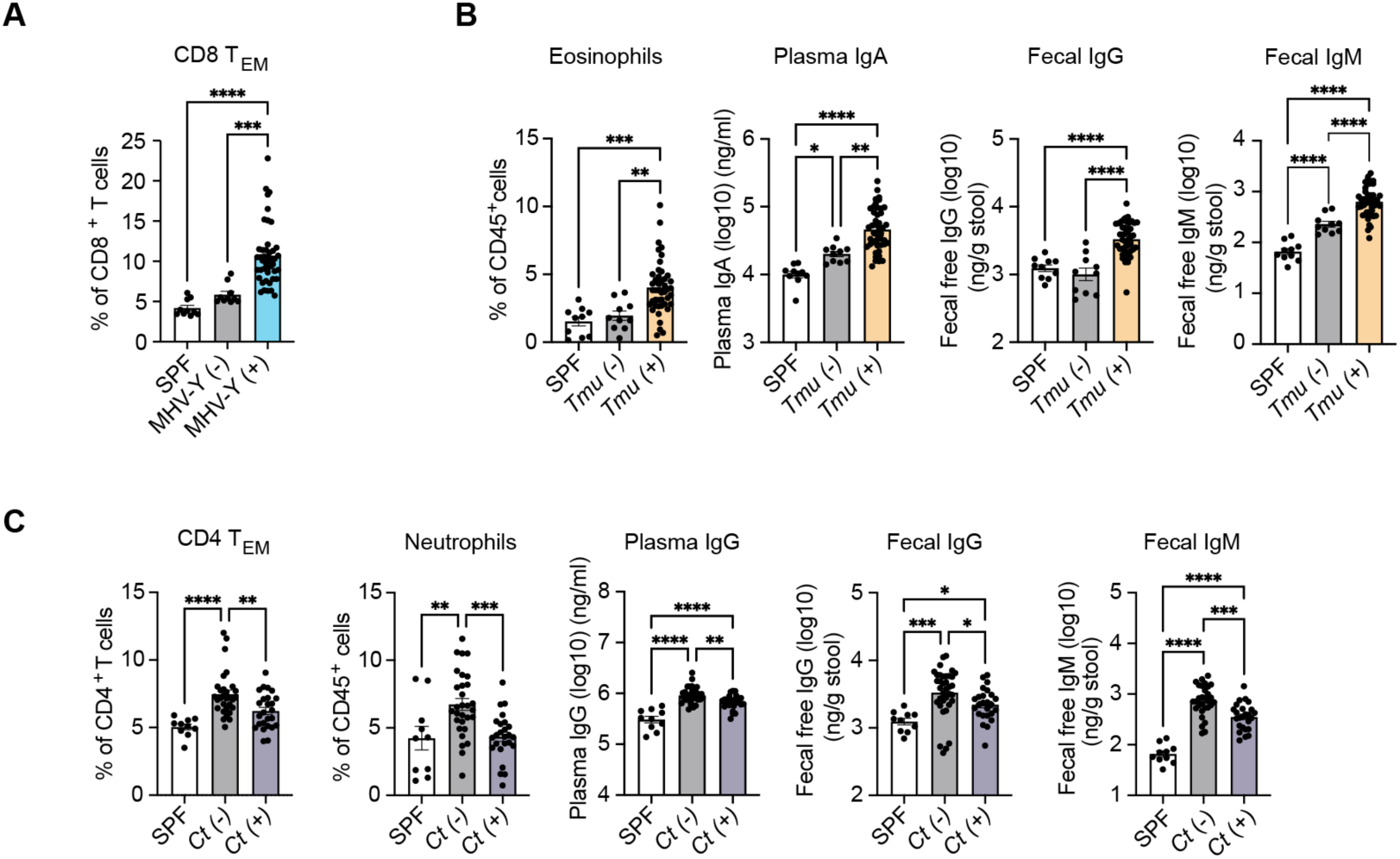
Distinct microbes are associated with specific immune phenotypes. Comparison of immune readouts across polymicrobial exposure conditions at 4 weeks post cohousing. (**A**) Blood CD62L^−^CD44^+^ CD8 T_EM_ cells in SPF mice (n = 10 mice), mice not exposed to MHV-Y (n = 10 mice), and mice exposed to MHV-Y (n = 45 mice). (**B**) Blood eosinophils, plasma IgA, fecal IgG, and fecal IgM in SPF mice (n = 10 mice), mice not exposed to *Tmu* (n = 10 mice), and mice exposed to *Tmu* (n = 45 mice). (**C**) Blood CD62L^−^CD44^+^ CD4 T_EM_ cells and neutrophils, plasma IgG, fecal IgG, and fecal IgM in SPF mice (n = 10 mice), mice not exposed to *Ct* (n = 30 mice), and mice exposed to *Ct* (n = 25 mice). Data are presented as mean ± SEM. Statistical analysis was performed using one-way ANOVA with Tukey post hoc test. \**P* < 0.05; \*\**P* < 0.01; \*\*\**P* < 0.001; \*\*\*\**P* < 0.0001.

**Fig. S5.**
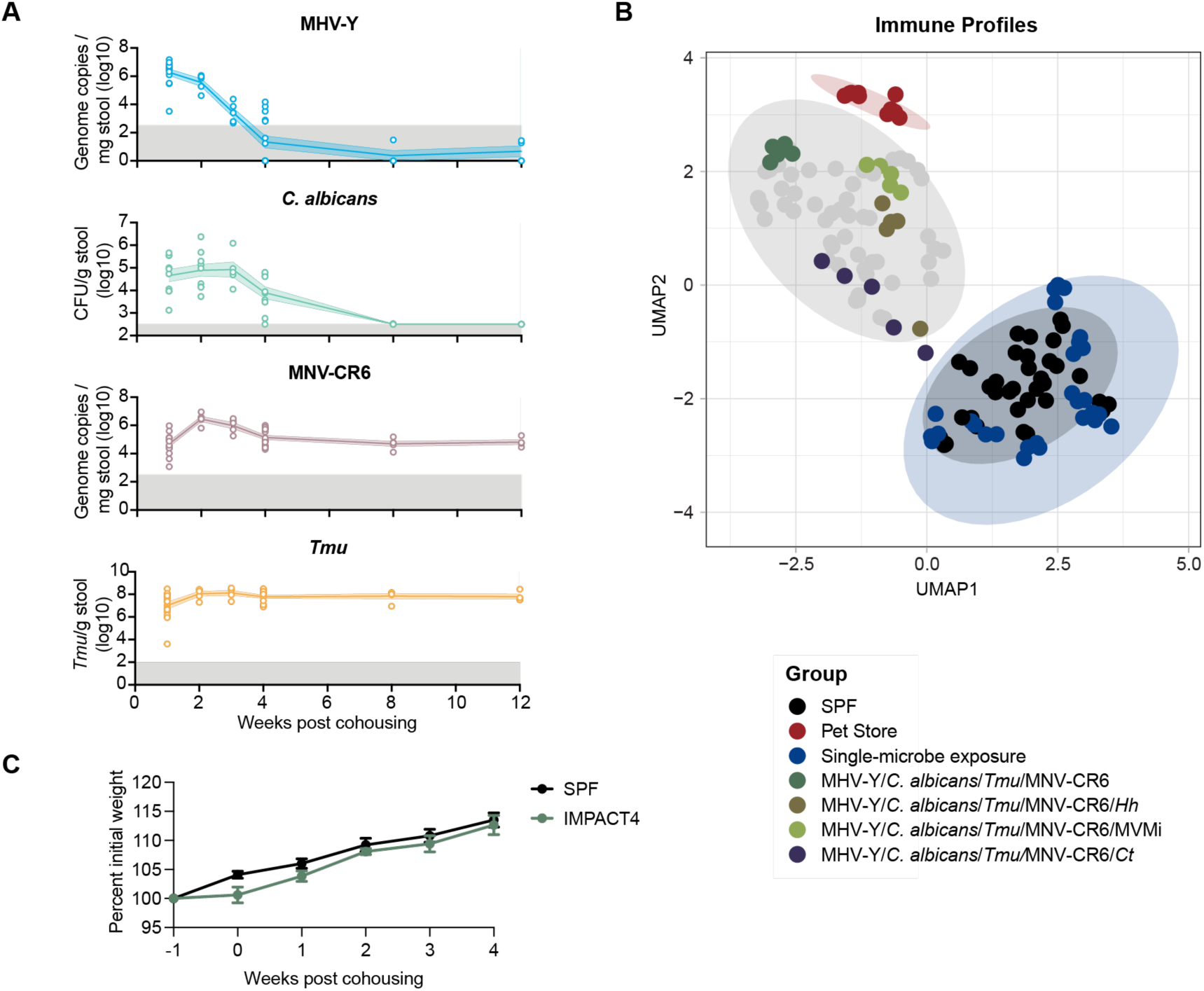
Optimization and characterization of the IMPACT4 consortium. (**A**) Longitudinal fecal burdens of individual microbes in mice exposed to IMPACT4. Gray shading indicates the limit of detection. (**B**) UMAP of immune profiles across microbial exposure conditions, highlighting mice exposed to IMPACT4 (MHV-Y/*C. albicans*/*Tmu*/MNV-CR6) or IMPACT4 with the addition of *Hh*, MVMi, or *Ct*. (**C**) Body weight of SPF (n = 5 mice) and IMPACT4 (n = 6 mice) mice. Statistical analysis was performed using two-way repeated-measures ANOVA (C). Data are pooled from two independent experiments (A) or representative of two independent experiments (C). Data are presented as mean ± SEM (A and C).

**Fig. S6.**
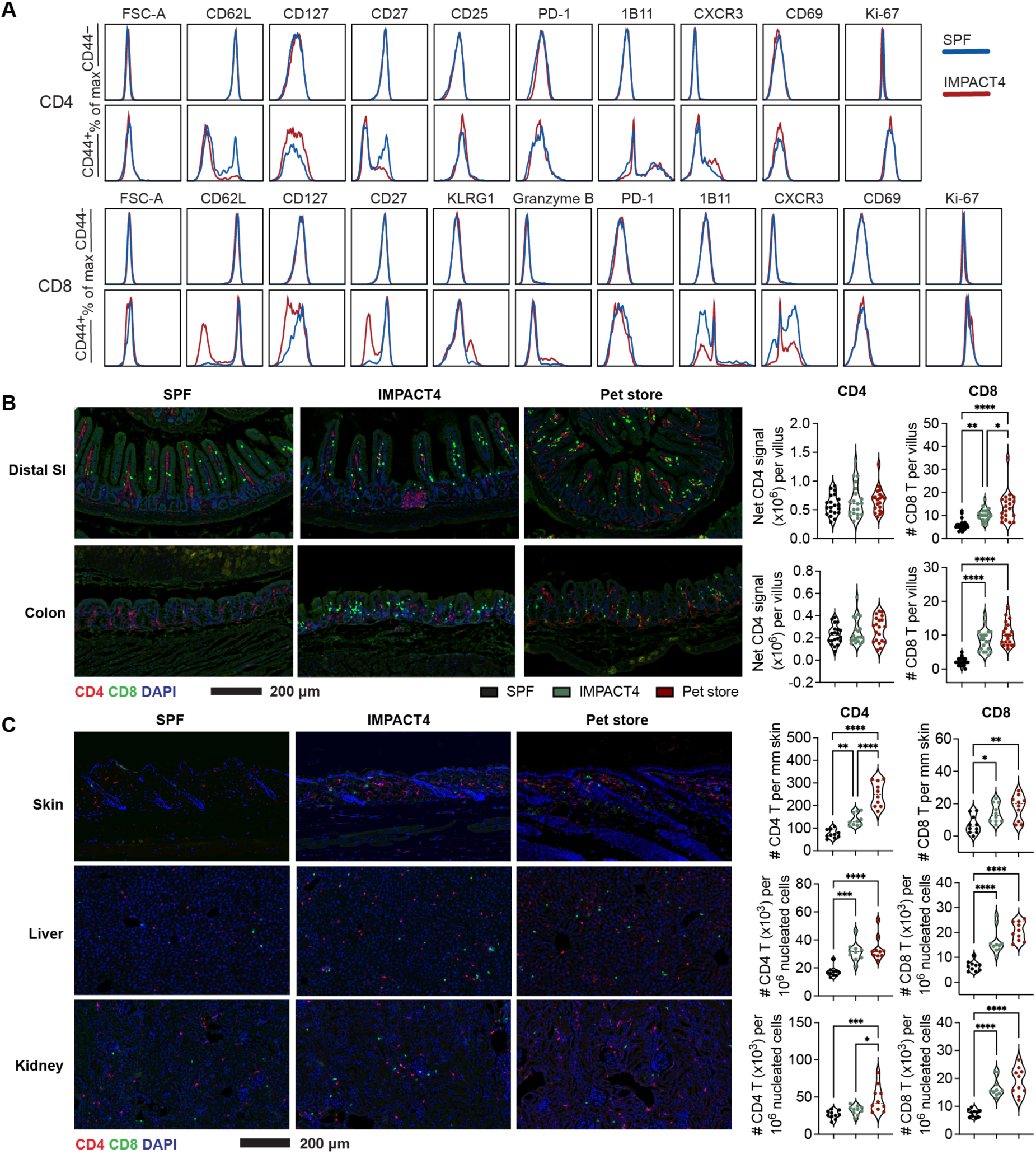
IMPACT4 induces systemic T cell remodeling. Samples were collected 4 weeks after IMPACT4 exposure. (**A**) Representative flow cytometry histograms of indicated surface markers on CD62L^+^CD44^−^ and CD44^+^ blood T cells from SPF and IMPACT4 mice. (**B** and **C**) Immunofluorescence staining and quantification of CD4 and CD8 T cells in distal small intestine (SI) and colon (B) and skin, liver, and kidney (C) from SPF, IMPACT4, and pet store mice. T cell density was quantified from 20 intestinal villi or 10 randomly selected fields of view using at least two mice per group. Representative images are shown. Statistical analysis was performed using one-way ANOVA with Tukey post hoc test (B and C). Data are representative of two independent experiments. Violin plots indicate median and quartiles (B and C). \**P* < 0.05; \*\**P* < 0.01; \*\*\**P* < 0.001; \*\*\*\**P* < 0.0001.

**Fig. S7.**
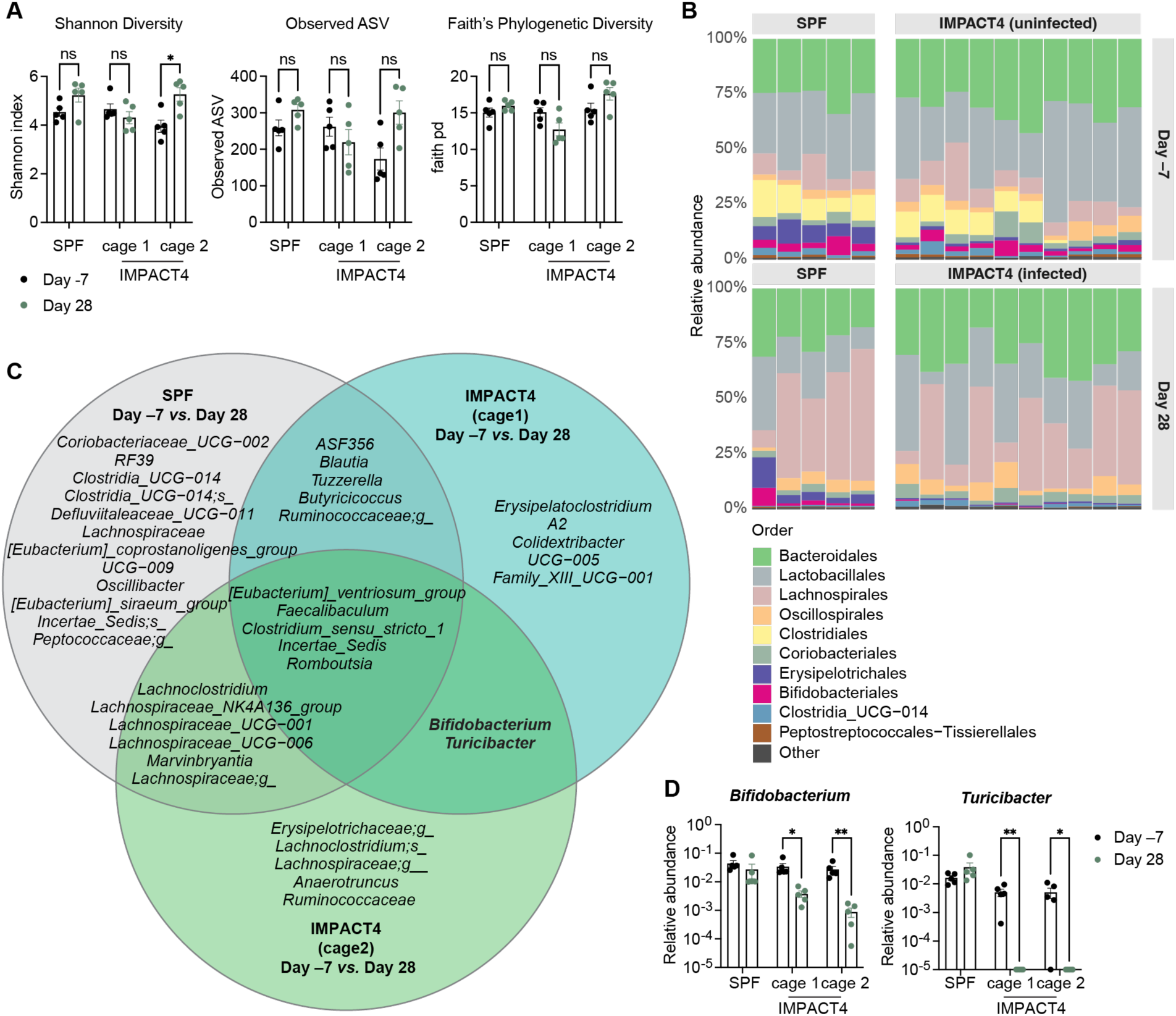
IMPACT4 minimally alters the resident microbiota. (**A**) Shannon diversity, observed ASVs, and Faith’s phylogenetic diversity of fecal microbiota at day −7 and day 28 across three independent cages. Day 28 corresponds to 28 days after microbial exposure for IMPACT4 cages. n = 5 mice per cage. (**B**) Relative abundance of bacterial taxa at the order level. Each bar represents one mouse. (**C**) Venn diagram of significantly altered bacterial taxa between day −7 and day 28 in SPF and IMPACT4 cages. Differential abundance analysis was performed separately within each cage using DESeq2. Taxa with adjusted P < 0.05 were included. (**D**) Relative abundance of *Bifidobacterium* and *Turicibacter* in SPF and IMPACT4 cages. n = 5 mice per cage. Statistical analysis was performed using multiple unpaired t tests (A and D). Data are presented as mean ± SEM (A and D). \**P* < 0.05; \*\**P* < 0.01; ns, not significant.

**Fig. S8.**
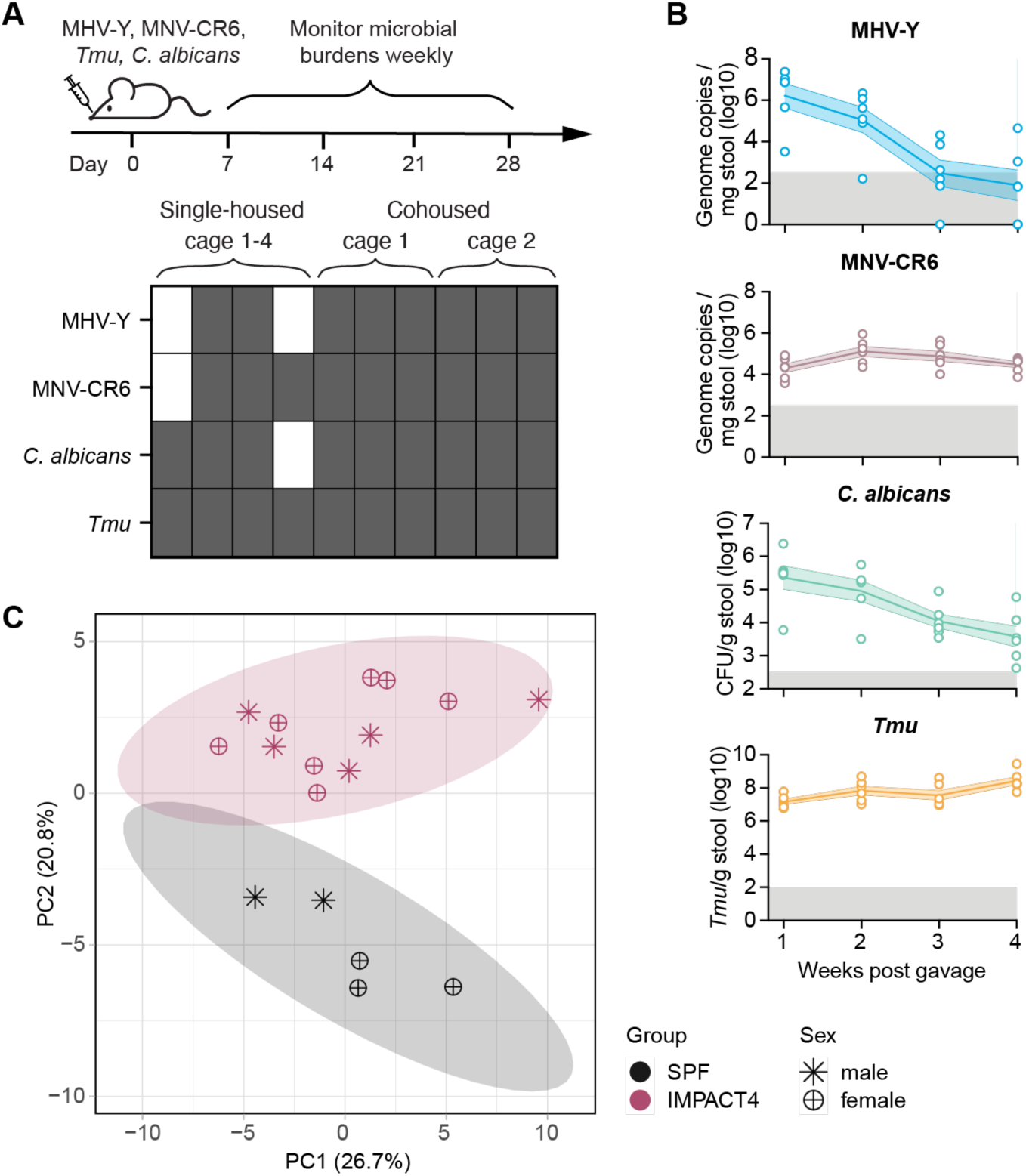
**Single-step oral gavage extends IMPACT4 to both sexes**. (**A**) Detection of IMPACT4 microbes in fecal samples 7 days after single-step oral gavage of the consortium. Gray indicates detection and white indicates no detectable signal. Columns represent individual mice annotated as single-housed or cohoused. (**B**) Longitudinal fecal burdens of individual microbes following gavage (n = 6 mice). Gray shading indicates the limit of detection. Data are representative of at least two independent experiments and are presented as mean ± SEM. (**C**) PCA of immune profiles in female and male mice before and after IMPACT4 exposure. Data are pooled from two independent experiments.

**Fig. S9.**
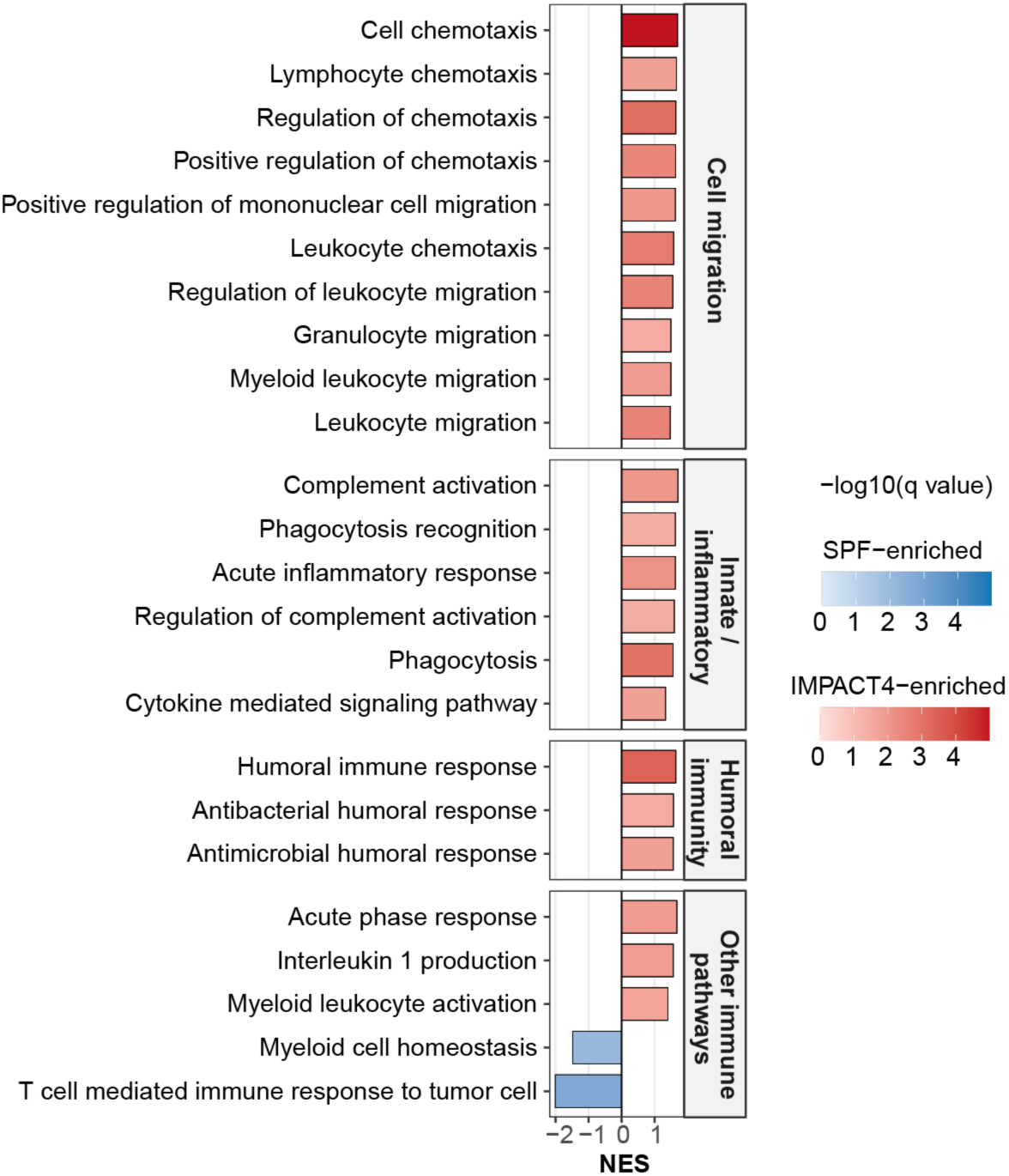
**IMPACT4 induces broad transcriptional remodeling of immune pathways in blood**. GSEA of Gene Ontology Biological Process pathways in blood from experiment-matched SPF and IMPACT4 mice collected 4 weeks after IMPACT4 exposure. Significantly enriched immune-related pathways are grouped by functional category and displayed by normalized enrichment score (NES). Bar color intensity indicates statistical significance [−log10(q value)]. Red denotes IMPACT4-enriched pathways and blue denotes SPF-enriched pathways. Only pathways with q < 0.05 are shown.

**Fig. S10.**
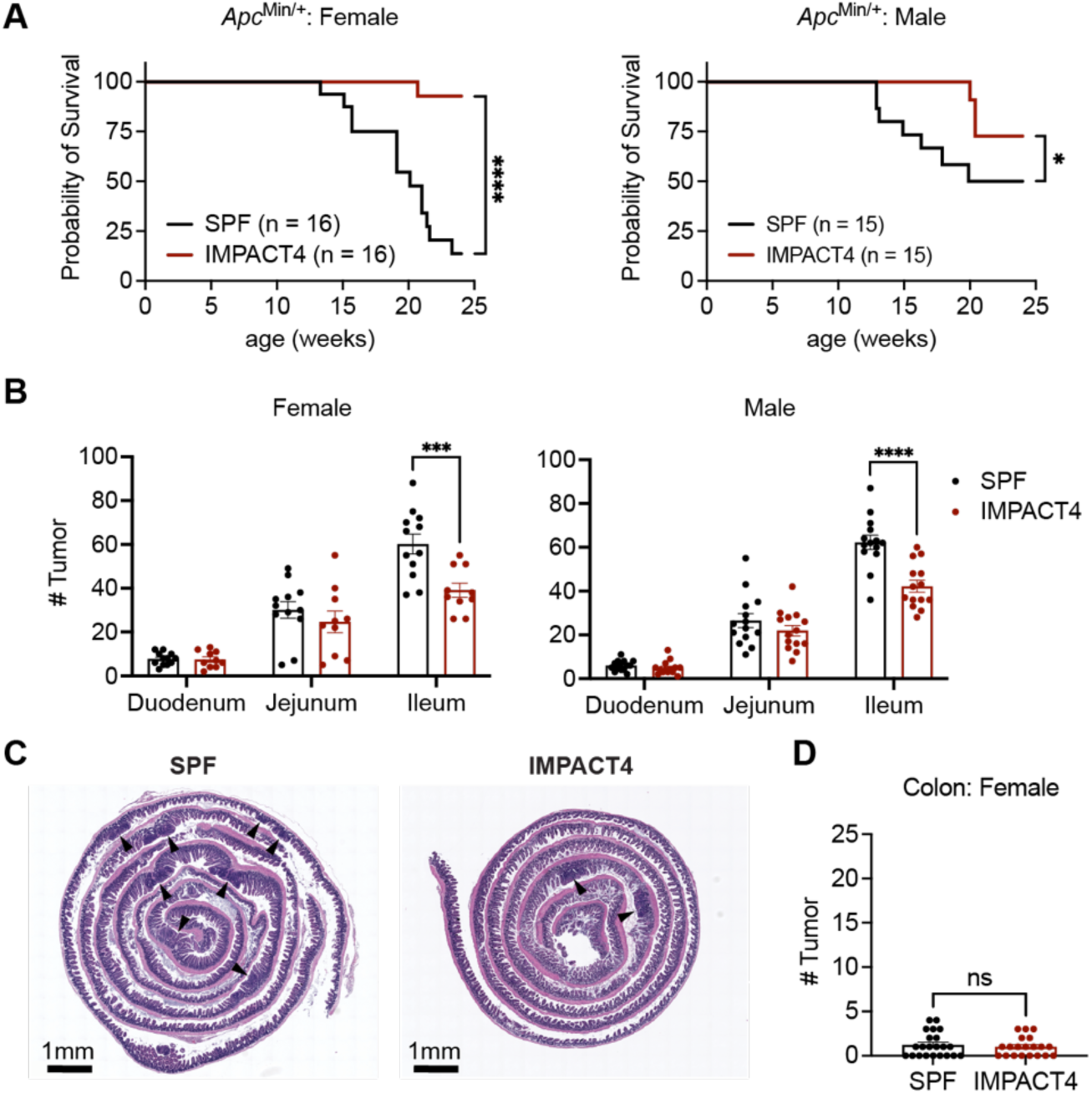
IMPACT4 alters tumorigenesis in *Apc*^Min/+^ mice. (**A**) Survival of female (left) and male (right) *Apc*^Min/+^ mice maintained under SPF conditions or exposed to IMPACT4 at 6 weeks of age. n = 16 female mice per group; n = 15 male mice per group. (**B**) Polyp numbers in intestinal segments of 16-week-old female (left) and male (right) *Apc*^Min/+^ mice. n = 12 SPF and 10 IMPACT4 female mice; n = 14 male mice per group. (**C**) Representative H&E-stained ileal Swiss rolls from 10-week-old *Apc*^Min/+^ mice. Arrowheads indicate polyps. (**D**) Colonic tumor numbers in female *Apc*^Min/+^ mice at 10 to 20 weeks of age. n = 22 SPF and 20 IMPACT4 mice. Statistical analysis was performed using log-rank Mantel-Cox test (A), two-way ANOVA with Sidak correction (B), or unpaired t test (D). Data are pooled from at least two independent experiments. Data are presented as mean ± SEM (B and D). \**P* < 0.05; \*\**P* < 0.01; \*\*\**P* < 0.001; \*\*\*\**P* < 0.0001; ns, not significant.

**Fig. S11.**
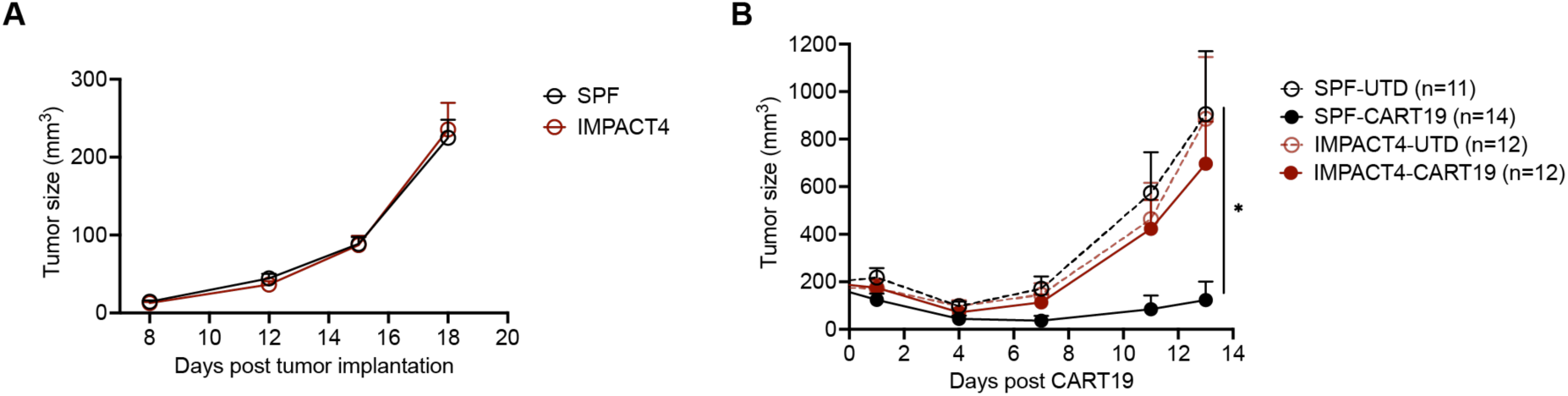
**IMPACT4 alters tumor response to CART19 treatment**. (**A**) A20 tumor growth before CART19 treatment in SPF and IMPACT4 BALB/c mice. n = 31 SPF and 29 IMPACT4 mice. (**B**) Tumor growth following UTD or CART19 transfer. n = 11 SPF and 12 IMPACT4 mice for UTD treatment; n = 14 SPF and 12 IMPACT4 mice for CART19 treatment. Statistical analysis was performed using two-way repeated-measures ANOVA with Sidak correction (A) or Tukey post hoc test (B). Data are pooled from two independent experiments and are presented as mean ± SEM. \**P* < 0.05.

**Fig. S12.**
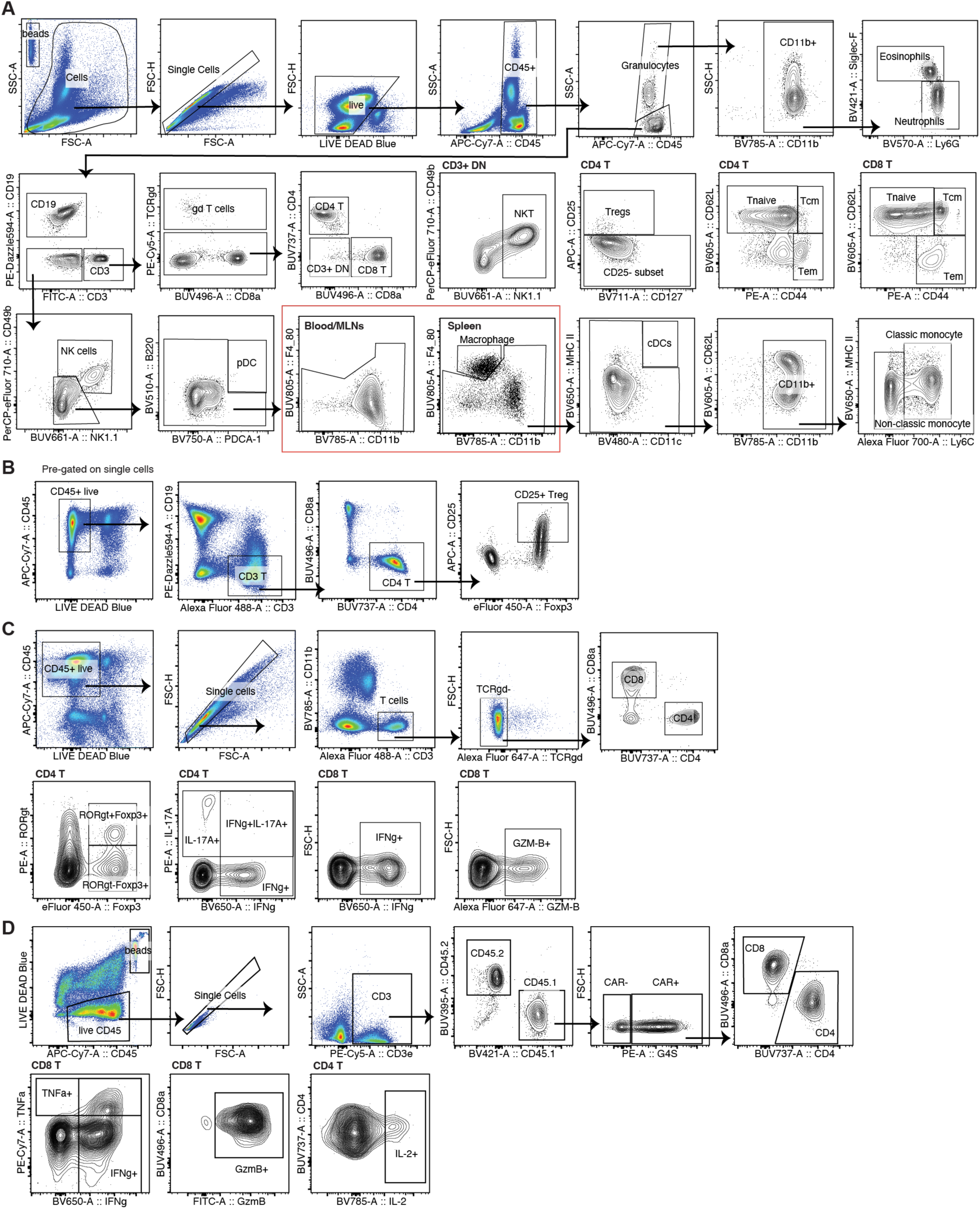
Flow cytometry gating strategies. (**A**) Gating strategy for major immune cell populations in blood, spleen, and mesenteric lymph nodes (MLN). A representative blood sample is shown. (**B**) Gating strategy for CD25^+^Foxp3^+^ Tregs in the CD28SA study. (**C**) Gating strategy for intestinal immune profiling in the ICB study. (**D**) Gating strategy for CAR T cells in spleen and tumor in the CART19 study. A representative spleen sample is shown.

**Table S1.** Microbial composition of tested combination in the screening.

| **Combinations** | **Microbes** |
| --- | --- |
| Combo 1 | MNV-CR6, MVMi, MHV-Y, *Hh*, *C. albicans* CHN1 |
| Combo 2 | MNV-CR6, MHV-Y, *Hh*, *C. albicans* CHN1, *Tmu* |
| Combo 3 | MNV-CR6, MVMi, *Hh*, *C. albicans* CHN1, *Tmu* |
| Combo 4 | MNV-CR6, MVMi, MHV-Y, *C. albicans* CHN1, *Tmu* |
| Combo 5 | MNV-CR6, MVMi, MHV-Y, *Hh*, *Tmu* |
| Combo 6 | MVMi, MHV-Y, *Hh*, *C. albicans* CHN1, *Tmu* |
| Combo 7 | MHV-Y, *Hh*, *C. albicans* CHN1, *Tmu*, *C. tyzzeri* STL |
| Combo 8 | MNV-CR6, *Hh*, *C. albicans* CHN1, *Tmu*, *C. tyzzeri* STL |
| Combo 9 | MNV-CR6, MHV-Y, *C. albicans* CHN1, *Tmu*, *C. tyzzeri* STL |
| Combo 10 | MNV-CR6, MHV-Y, *Hh*, *Tmu*, *C. tyzzeri* STL |
| Combo 11 | MNV-CR6, MHV-Y, *Hh*, *C. albicans* CHN1, *C. tyzzeri* STL |

**Table S2.**
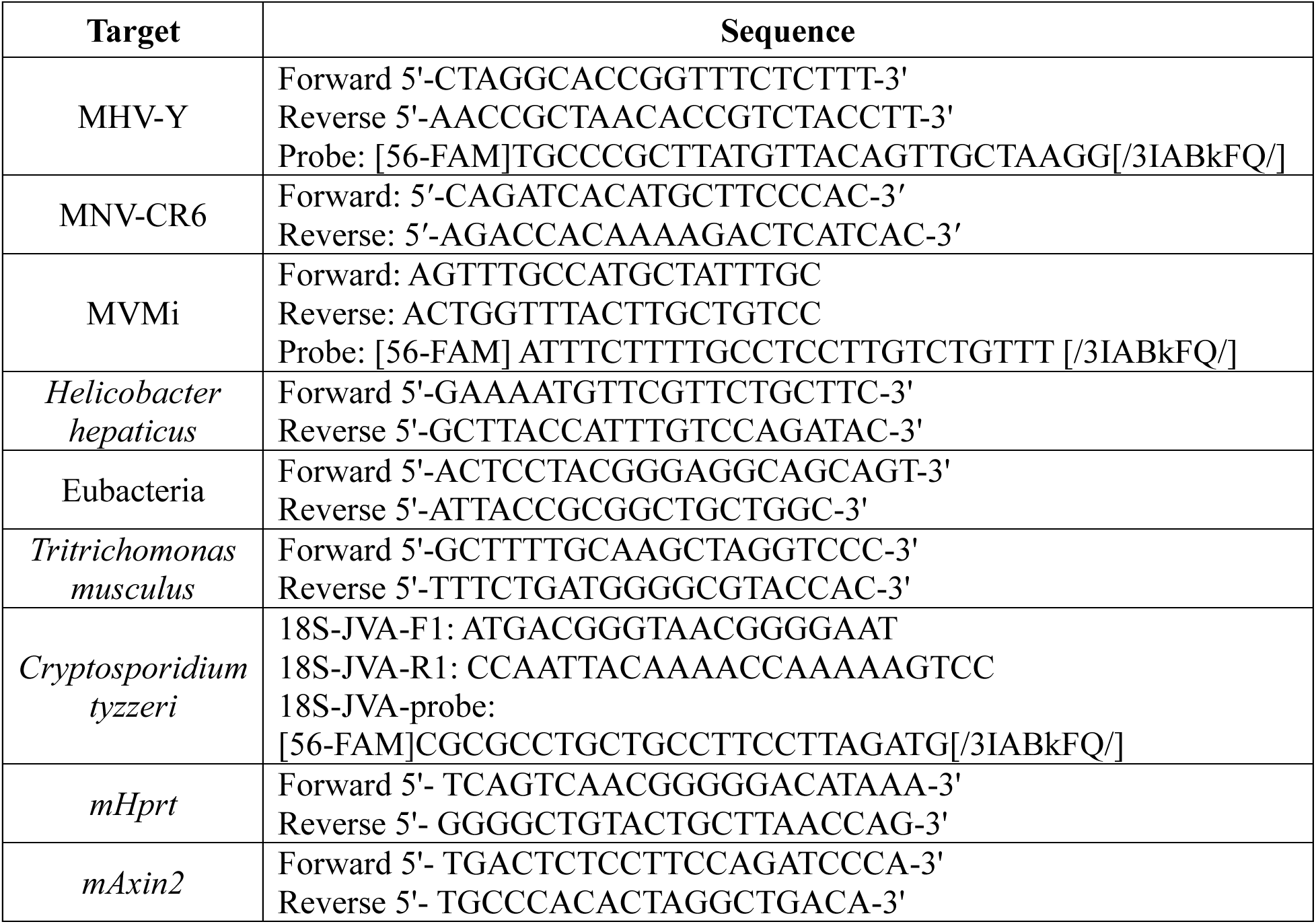
Primers and probes used in this study.

**Table S3.**
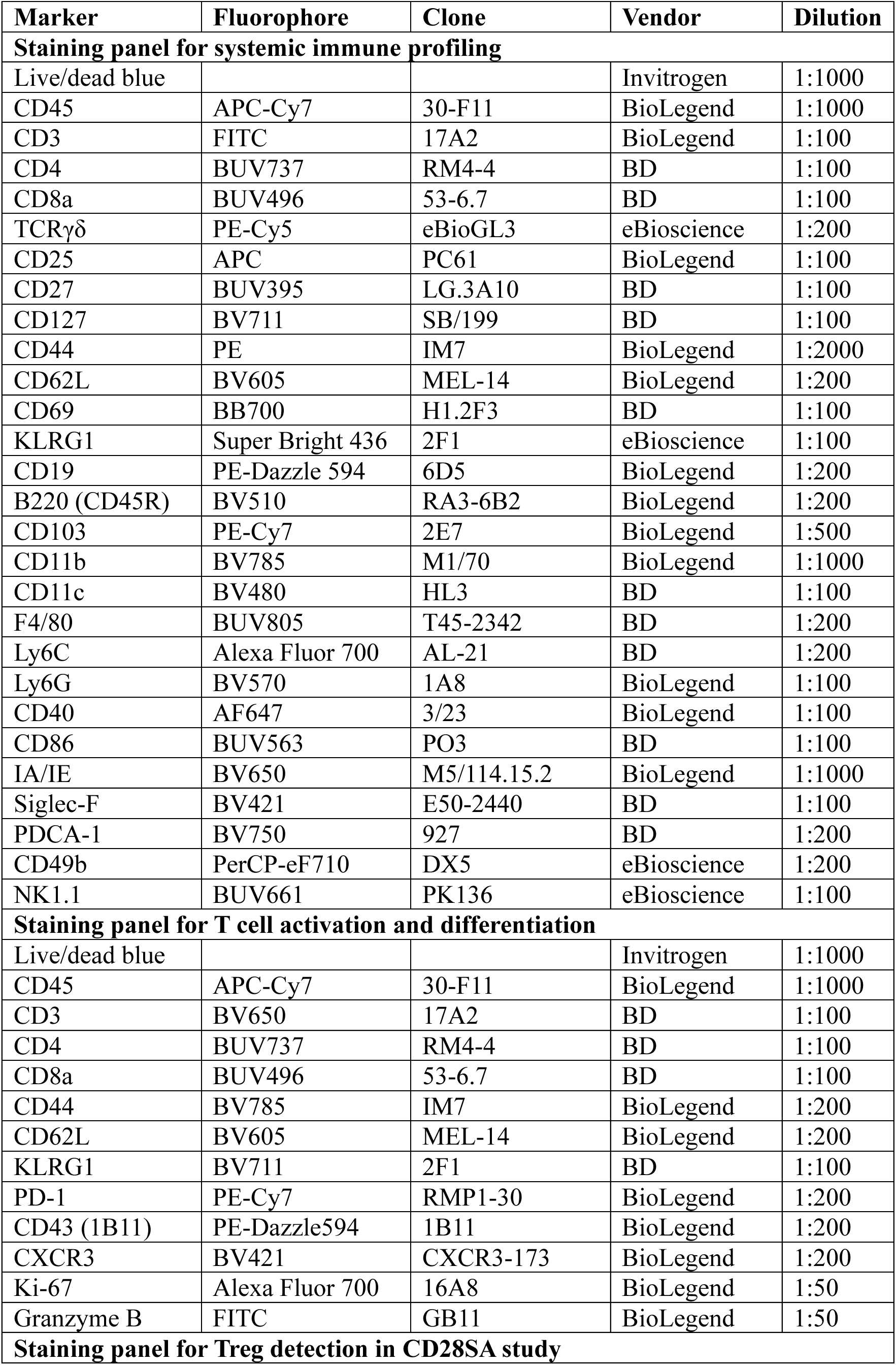
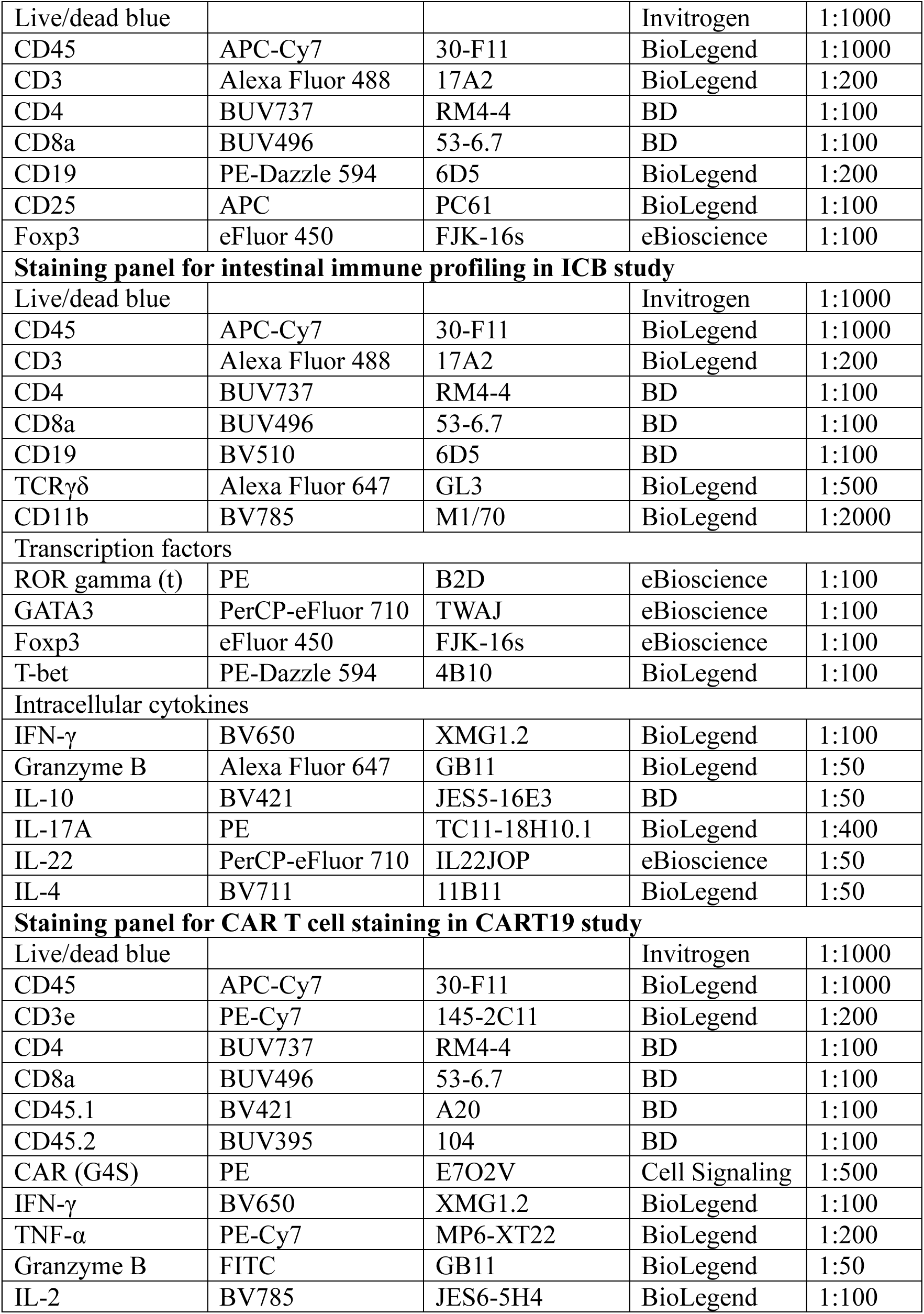
Flow cytometry staining panels used in this study.

## Materials and Methods

### Mice

Wild-type C57BL/6 and BALB/c mice were obtained from The Jackson Laboratory, Charles River Laboratories, Taconic Biosciences, or generated from mice bred onsite in the animal facility of the Vernon and Shirley Hill Pavilion at the University of Pennsylvania. Mice obtained from external vendors were acclimated to the animal facility for at least 1 week before experimental use. C57BL/6J-*Apc*^Min/+^/J mice were originally obtained from The Jackson Laboratory (strain no. 002020) and subsequently bred in the animal facility at the Vernon and Shirley Hill Pavilion.

Mice were maintained under specific pathogen-free (SPF) conditions in individually ventilated cages on a 12-hour light/dark cycle with ad libitum access to irradiated chow and sterile water. Female mice were used for experiments involving cohousing, whereas both male and female mice were used in studies that did not require cohousing. Mice were age-and sex-matched within each experiment.

Germ-free C57BL/6 mice for generating protist stocks were bred in flexible-film isolators in the University of Pennsylvania Gnotobiotic and Naturalized Mouse Core. For experiments, germ-free mice were transferred to ISOcages (Tecniplast) with sterile food and water, and absence of bacteria was monitored throughout the experiment by qPCR for 16S rRNA gene. Pet store mice were purchased from local pet stores in the Philadelphia area. The genetic backgrounds of pet store mice were assessed using the MiniMUGA genotyping panel (v2.3.1; Transnetyx, Cordova, TN, USA).

All animal experiments were conducted in accordance with protocols approved by the Institutional Animal Care and Use Committee (IACUC) of the University of Pennsylvania.

### Cell lines

293T cells (ATCC), BHK-mCEACAM1 cells (from Dr. Siddharth Krishnamurthy) and BV2 cells (a gift from Dr. Timothy Nice) were cultured in DMEM (Corning) supplemented with 50 U/mL penicillin and 50 mg/mL streptomycin (Corning), plus 10% heat-inactivated fetal bovine serum (FBS) (Cytiva). NB324K cells (a gift from Dr. Peter Tattersall) were cultured in DMEM supplemented with 50 U/mL penicillin and 50 mg/mL streptomycin plus 5% heat-inactivated FBS. A20 cells (ATCC) were cultured in ‘‘R10’’ (RPMI 1640 + 10% FBS, 50 U/mL penicillin and 50 mg/mL streptomycin, 10 mM HEPES, 1% GlutaMAX) medium supplemented with 0.05 mM 2-mercaptoethanol. All cell lines are cultured at 37°C in 5% ambient CO_2_.

### Microbial strains

#### MNV-CR6

Molecular clones of the MNV strain CR6 were generated as previously described to produce working stocks of infectious virus (*45*). Plasmids encoding the viral genome were transfected into 293T cells to generate a P0 stock. This P0 stock was subsequently amplified through two passages in BV2 cells at a MOI of 0.05 to produce P2 stock. For stock preparation, infected BV2 cells and culture supernatants were collected, freeze-thawed, centrifuged at 1,200 × g for 5 min, filtered through a 0.22-μm filter, and concentrated using 100-kDa MWCO Amicon Ultra centrifugal filters. Viral titers were determined by plaque assay using BV2 cells overlaid with DMEM (Corning) containing 1% methylcellulose (Sigma-Aldrich). Plaques were visualized by crystal violet staining 3 days post infection. Mice were infected by oral gavage with 3 × 10^6^ PFU of MNV-CR6.

#### MHV-Y

The P1 stock of MHV-Y was produced and provided by Dr. Siddharth Krishnamurthy. The P1 stock was amplified in BHK-mCEACAM1 cells at a multiplicity of infection (MOI) of 0.01–0.1 in the Cadwell lab to generate P2 working stocks used for mouse infection. For virus propagation, infected cultures were subjected to freeze-thaw lysis and centrifuged at 3,000 rpm for 10 min at 4°C. Supernatants were collected and stored as viral stocks. Viral titers were determined by the tissue culture infectious dose 50 (TCID50) assay. Briefly, 2 × 10^4^ BHK-mCEACAM1 cells were seeded per well in 96-well plates. Viral stocks were serially diluted 10-fold and added to cells. Plates were incubated at 37°C for 48 h, and viral titers were determined based on the presence of syncytia. TCID50 values were calculated using the Reed–Muench method with 10 replicate wells per dilution. Mice were infected by oral gavage with 1 × 10^4^ TCID50 of MHV-Y.

#### MVMi

MVMi stocks were originally provided by Dr. David Pintel (University of Missouri, Columbia, MO, USA). MVMi propagation and titer determination were performed as previously described (*17*). Briefly, viral stocks were expanded in NB324K cells, and culture supernatants were collected and stored as virus stocks. Viral titers were determined by focus-forming assay on NB324K cells. For titer determination, viral stocks were serially diluted in DMEM and added to nearly confluent NB324K cells seeded in 96-well clear-bottom black plates (Corning). Cells were incubated at 37°C for 1 h with gentle agitation every 15 min. The inoculum was then removed and replaced with DMEM supplemented with 10% FBS. At 24 h post infection, cells were fixed in HyClone Water (GE Healthcare Life Sciences) containing 2% paraformaldehyde for 20 min on ice and permeabilized in quench/permeabilization buffer (20 mM glycine and 0.25% Triton X-100 in PBS) for 20 min on ice. MVM-infected cells were detected using a non-commercial anti-MVM NS protein antibody provided by Dr. Peter Tattersall. Following incubation with the primary antibody for 1 h on ice, cells were stained for 15 min on ice with Alexa Fluor 488-conjugated anti-mouse IgG secondary antibody (Thermo Fisher Scientific) and DAPI (Sigma-Aldrich). Plates were imaged using an EVOS Cell Imaging System (Thermo Fisher Scientific), and focus-forming units (FFU) were quantified manually using ImageJ (NIH). Mice were infected by oral gavage with 2 × 10^5^ FFU of MVMi.

#### C. albicans CHN1

A single colony of *Candida albicans* CHN1 was cultured in 5 ml of Sabouraud dextrose media (BD Difco) at 30°C for approximately 16 h with shaking. Cells were pelleted by centrifugation at 1,000 × g for 5 min and resuspended in 1.5 mL of PBS. Mice were orally gavaged with 100 μL of *C. albicans* [approximately 2 × 10^7^ colony-forming units (CFUs) per mouse].

### Tritrichomonas musculus

Purified *Tritrichomonas musculus* (*Tmu*) protists were originally provided by Dr. Michael R. Howitt (Stanford University). *Tmu* was expanded in germ-free C57BL/6 mice. Cecal contents were collected 2 weeks after inoculation, passed through a 40-μm cell strainer, and washed three times with PBS. Protists were then purified from the interface of a 40%/80% Percoll (Sigma-Aldrich) gradient following centrifugation at 1,000 × g for 10 min with no brake. Viable protists were enumerated using a hemocytometer. For cryopreservation, protists were pelleted at 700 × g for 5 min and resuspended in half the final volume of complete PBF medium prepared as previously described (*16*). An equal volume of 2× freezing medium (complete PBF medium containing 15% DMSO) was then added to achieve a final DMSO concentration of 7.5%. Protists were aliquoted into cryovials and immediately placed at −80°C in a Mr. Frosty freezing container. Cryovials were transferred to liquid nitrogen storage 8–24 h later.

For mouse colonization, frozen *Tmu* stocks were thawed rapidly in a 37°C water bath and diluted into 9 mL PBS. Protists were pelleted by centrifugation at 100 × g for 5 min, resuspended in PBS, and administered by oral gavage. Each mouse was infected by oral gavage of 1 × 10^5^ viable protists.

### Cryptosporidium tyzzeri STL

*Cryptosporidium tyzzeri* (*Ct*) STL oocysts were originally provided by Dr. Chyi-Song Hsieh (Washington University School of Medicine) (*20*). Transgenic strain expressing NanoLuc luciferase and tdTomato were provided by Dr. Boris Striepen (University of Pennsylvania) (*46*). Parasites were propagated in *Ifng*^-/-^ mice (The Jackson Laboratory, stock no. 002287). Oocysts were purified from fecal samples by sucrose flotation followed by cesium chloride gradient centrifugation as previously described (*47*). Briefly, fecal pellets were homogenized in tap water using a LabGen 125 homogenizer (Cole-Parmer, Vernon Hills, IL, USA) and passed through a 250-μm mesh filter. The filtrate was suspended in tap water and centrifuged at 1,000 × g for 5 min. The pellet was resuspended in tap water and was mixed 1:1 with saturated sucrose solution (specific gravity 1.33) and centrifuged at 1,000 × g for 5 min. The resulting supernatant was collected, diluted in 0.85% saline, overlaid onto a cesium chloride solution (specific gravity 1.15), and centrifuged at 16,000 × g for 3 min. Purified oocysts were collected from the saline-CsCl interface and resuspended in cold PBS. Mice were infected by oral gavage with 1 × 10^4^ oocysts per mouse.

### Helicobacter hepaticus

*H. hepaticus* stocks were provided by Dr. Dan Littman (New York University). Frozen stock aliquots were maintained in Brucella broth supplemented with 20% glycerol at −80°C. For bacterial expansion, *H. hepaticus* was streaked onto TSA plates containing 5% sheep blood (Thermo Fisher Scientific). Inoculated plates were placed in AnaeroPack Rectangular Jars (Mitsubishi Gas Chemical) containing sterile water to maintain humidity. Jars were equilibrated for 30 min in an anaerobic chamber supplied with 90% nitrogen and 10% carbon dioxide (Airgas), sealed, and transferred to a 37°C incubator. Under these conditions, bacteria were cultured in a humid microaerophilic environment for 5 days prior to animal inoculation. For oral infection, bacteria were collected from agar plates and resuspended in Brucella broth. Bacterial density was estimated by optical density at 600 nm (OD600) and adjusted to an OD600 of 2.0. Each mouse received 100 μL of bacterial suspension by oral gavage.

### Microbial burden quantification

Fresh fecal pellets were collected and maintained on ice prior to processing. To enable quantification of multiple microbes from the same sample, fecal pellets were weighed, resuspended in 600 μL sterile PBS, and homogenized. For nucleic acid extraction, 400 μL of homogenate was mixed with an equal volume of DNA/RNA Shield 2× Concentrate (Zymo Research). The remaining homogenate was used for quantification of *C. albicans*.

For fungal burden measurements, fecal homogenates were serially diluted 10-fold in sterile PBS and spot-plated onto Sabouraud dextrose agar supplemented with chloramphenicol (25 μg/mL; Sigma-Aldrich). Colonies were enumerated following overnight incubation at 30°C, and CFU were normalized to fecal weight.

For all other microbes, burden was quantified by qPCR. DNA and RNA were isolated from the same fecal sample using the ZymoBIOMICS DNA/RNA Mini Kit (Zymo Research) according to the manufacturer’s instructions. DNA and RNA were eluted in 100 μL and 50 μL nuclease-free water, respectively. For MHV-Y and MNV-CR6, RNA was reverse transcribed using the High-Capacity cDNA Reverse Transcription Kit (Applied Biosystems), and resulting cDNA was used for qPCR analysis. For MVMi, *Tmu*, *Ct*, and *Hh*, diluted DNA was used directly as template.

TaqMan Universal PCR Master Mix (Applied Biosystems) was used for detection of MHV-Y, MVMi, and *Ct*, whereas PowerTrack SYBR Green Master Mix (Applied Biosystems) was used for MNV-CR6, *Hh*, and *Tmu*. Reactions were performed on a QuantStudio 5 Real-Time PCR System (Applied Biosystems). For *Hh*, microbial burden was calculated by normalizing *Hh* Ct values to total bacterial abundance measured using universal eubacterial primers. For all other microbes, microbial burden was quantified using standard curves and normalized to fecal weight. Standard curves were generated using linearized plasmid standards for MHV-Y, MNV-CR6, and MVMi, or nucleic acids extracted from a known number of *Tmu* cells or *Ct* oocysts.

Primer and probe sequences are listed in table S2.

### Polymicrobial infection, immune profiling, and immunoglobulin quantification

Six-week-old mice were exposed to the IMPACT4 consortium either through cohousing or a single-step infection strategy. For the cohousing approach, mice were first infected individually with one microbe by oral gavage and housed separately. One week later, after successful infection had been confirmed by fecal microbial quantification, mice carrying different microbes were cohoused. For the single-step infection strategy, mice were simultaneously gavaged with MHV-Y, MNV-CR6, *C. albicans* CHN1 and *Tmu*. Successful polymicrobial exposure was confirmed by fecal microbial burden measurements one week after completion of the infection protocol.

For systemic immune profiling, tissues were collected 4 weeks after microbial exposure. Blood was collected by cardiac puncture, and plasma was isolated by centrifugation at 2,000 rpm for 5 min at 4°C, snap-frozen, and stored at −80°C. Spleens and mesenteric lymph nodes (MLN) were passed through 70-μm cell strainers and resuspended in FACS buffer (PBS containing 1% BSA, 1 mM EDTA, 20 mM HEPES, and 1 mM sodium pyruvate). Red blood cells from blood and spleen samples were lysed using ACK lysis buffer (Gibco) prior to staining. For flow cytometric analysis, 100 μL whole blood, 1/20 of splenocytes, and 1/3 of MLN cells were stained. Cells were stained with LIVE/DEAD Blue Fixable Viability Dye (Invitrogen), followed by Fc receptor blockade using TruStain FcX™ PLUS (BioLegend). Surface staining was performed using antibodies against CD45, CD3, CD4, CD8a, TCRγδ, CD25, CD27, CD127, CD44, CD62L, CD69, KLRG1, CD19, B220, CD103, CD11b, CD11c, F4/80, Ly6C, Ly6G, CD40, CD86, IA/IE, Siglec-F, PDCA-1, CD49b, and NK1.1 (table S3). A separate flow cytometry panel was used to characterize T cell activation and differentiation in blood from IMPACT4-exposed mice (table S3). Surface staining was performed using antibodies against CD45, CD3, CD4, CD8a, CD44, CD62L, KLRG1, PD-1, 1B11, and CXCR3. Intracellular staining for Ki-67 and granzyme B was performed using the Foxp3/Transcription Factor Staining Buffer Set (eBioscience) according to the manufacturer’s instructions. Precision Count Beads (BioLegend) were added for absolute cell quantification. Samples were acquired on an Aurora flow cytometer (Cytek Biosciences) and analyzed using FlowJo (version 10.10.0). Gating strategies are shown in fig. S12A.

Total IgG, IgA, IgM, and IgE levels in plasma and fecal supernatants were quantified by ELISA. High-binding 96-well plates (Corning) were coated overnight at 4°C with capture antibodies diluted in PBS. Capture antibodies were used at 2 μg/mL and included goat anti-mouse IgG Fc-specific antibody (Sigma-Aldrich, Cat. #M2650), rat anti-mouse IgE antibody (BD Biosciences, Cat. #553413), goat anti-mouse IgA α-chain-specific antibody (Sigma-Aldrich, Cat. #M8769), and rat anti-mouse IgM antibody (BioLegend, Cat. #406502). Plates were washed with PBS containing 0.05% Tween-20 and blocked with PBS containing 2% BSA and 0.05% Tween-20 for 2 h at room temperature. Plasma or fecal supernatant samples were diluted in blocking buffer and incubated on coated plates for 2 h at room temperature. Standard curves were generated using mouse IgG isotype control (Invitrogen, Cat. #10400C), purified mouse IgE κ isotype standard (BD Biosciences, Cat. #553481), purified mouse IgA κ isotype control (BD Biosciences, Cat. #553476), or mouse IgM isotype control (Sigma-Aldrich, Cat. #M5909). Plates were washed and incubated for 1 h at room temperature with HRP-conjugated detection antibodies diluted in blocking buffer. Detection antibodies included goat anti-mouse IgG (H+L)-HRP (Invitrogen, Cat. #31430), goat anti-mouse IgE-HRP (Invitrogen, Cat. #SA5-10263), goat anti-mouse IgA α-chain-specific-HRP (Sigma-Aldrich, Cat. #A4789), and goat anti-mouse IgM μ-chain-specific-HRP (Jackson ImmunoResearch, Cat. #115-035-075). After washing, plates were developed with TMB substrate (eBioscience) and stopped with sulfuric acid. Absorbance was measured at 450 nm. Immunoglobulin concentrations were calculated from standard curves. Fecal immunoglobulin concentrations were normalized to fecal weight.

### CD28SA treatment and immune profiling

CD28SA monoclonal antibody (Bio X Cell, Cat. #BE0328) was administered intraperitoneally at 300 μg per mouse. Mice were euthanized either 1 day prior to treatment (baseline) or 96 h after CD28SA administration, and spleens were collected for immune profiling. Splenocytes were stained with LIVE/DEAD Blue Fixable Viability Dye (Invitrogen), followed by Fc receptor blockade using TruStain FcX™ PLUS (BioLegend). Cell surface staining was performed using antibodies against CD45, CD3, CD4, CD8a, CD19, and CD25 (table S3). Foxp3 staining was performed using the Foxp3/Transcription Factor Staining Buffer Set (eBioscience). Samples were acquired on an Aurora flow cytometer (Cytek Biosciences) and analyzed using FlowJo (version 10.10.0). Gating strategies are shown in fig. S12B.

For serum cytokine analysis, blood was collected by retro-orbital bleeding at baseline, 2 h, and 96 h following CD28SA injection. Serum was isolated, snap-frozen, and stored at −80°C until analysis. Cytokine concentrations were quantified using the LEGENDplex™ Mouse Inflammation Panel (13-plex) (BioLegend) according to the manufacturer’s instructions. Samples were acquired on an Aurora flow cytometer (Cytek Biosciences) and analyzed using the LEGENDplex™ data analysis software suite.

### Anti-CTLA-4 treatment, cecal immune profiling, and lipocalin-2 measurement

Four weeks after IMPACT4 exposure, mice received intraperitoneal injections of 200 μg anti-CTLA-4-mIgG2a antibody (clone 9D9, InvivoGen) or mouse IgG2a isotype control antibody (clone C1.18.4, Bio X Cell) every 3 days for a total of four doses. Fecal pellets were collected throughout the experiment, snap-frozen, and stored at −80°C until analysis. Mice were euthanized on day 12, and cecal tissues were harvested for immune profiling.

Cecal tissues were opened longitudinally, washed extensively with cold HBSS (Gibco), and cut into approximately 2-mm pieces. To remove epithelial cells, tissues were first incubated in buffered HBSS (HBSS supplemented with 20 mM HEPES and 1 mM sodium pyruvate) containing 5 mM EDTA and 1 mM DTT at 37°C with agitation for 15 min. Tissues were then vigorously shaken by hand for 2 min and passed through a metal strainer to remove released epithelial cells. Tissue fragments were subsequently incubated in buffered HBSS containing 1 mM DTT at 37°C with agitation for 10 min, followed by a second 2-min vigorous hand-shaking step. Tissues were again collected on a metal strainer, washed extensively with cold HBSS, minced, and digested in RPMI supplemented with 10% FBS, 10 mM HEPES, 0.5 mg/mL collagenase D (Sigma-Aldrich), and 0.01 mg/mL DNase I (Sigma-Aldrich) for 30 min at 37°C with agitation. Digested tissues were passed through a 70-μm cell strainer, and immune cells were enriched by Percoll gradient centrifugation using 40% and 80% Percoll (Sigma-Aldrich). Cells were collected from the 40%–80% interface following centrifugation at 900 × g for 22 min with no brake.

For intracellular cytokine analysis, half of the recovered cells were stimulated with Cell Stimulation Cocktail plus protein transport inhibitors (eBioscience) for 3.5 h at 37°C. Cells were stained using separate panels for transcription factors and intracellular cytokines. For both panels, cells were stained with LIVE/DEAD Blue Fixable Viability Dye (Invitrogen), followed by Fc receptor blockade using TruStain FcX™ PLUS (BioLegend). Surface staining was performed using antibodies against CD45, CD3, CD4, CD8a, CD19, TCRγδ, and CD11b (table S3). Transcription factor staining was performed using the Foxp3/Transcription Factor Staining Buffer Set (eBioscience). For intracellular cytokine staining, cells were fixed in Fixation Buffer (BioLegend) for 20 min at room temperature and stained with antibodies against IFN-γ, granzyme B, IL-10, IL-17A, IL-22, and IL-4 (table S3) using Intracellular Staining Perm Wash Buffer (BioLegend, Cat. No. 421002). Samples were acquired on an Aurora flow cytometer (Cytek Biosciences) and analyzed using FlowJo (version 10.10.0). Gating strategies are shown in fig. S12C.

For lipocalin-2 quantification, fecal pellets were weighed, homogenized in sterile PBS, and centrifuged at 1,000 × g for 10 min at 4°C. Lipocalin-2 concentrations in fecal supernatants were measured using the Mouse Lipocalin-2 DuoSet ELISA kit (R&D Systems) according to the manufacturer’s instructions.

### Intestinal tumor model in Apc^Min/+^ mice

C57BL/6J-*Apc*^Min/+^/J mice were maintained under SPF conditions or exposed to the IMPACT4 consortium at 6 weeks of age. To quantify intestinal tumors, mice were euthanized at 10 or 16 weeks of age. The intestine was dissected, measured, and the small intestine was divided into three equal-length segments corresponding to the duodenum, jejunum, and ileum. Intestinal segments were flushed with cold PBS, opened longitudinally, and laid flat on a dissection tray. Visible polyps were enumerated under a dissecting microscope, regardless of size. Spleens were collected and weighed at the 16-week time point. The ileum was prepared as a Swiss roll, fixed overnight in 4% paraformaldehyde, and transferred to 70% ethanol for histological analysis.

For gene expression analysis, two to three small tissue pieces were dissected from tumor-rich and tumor-free regions of the ileum from each 10-week-old mouse and stored in RNAlater (Invitrogen) until RNA extraction. RNA was extracted using the RNeasy Mini Kit (Qiagen) with on-column DNase digestion according to the manufacturer’s instructions. RNA was reverse-transcribed using the High-Capacity cDNA Reverse Transcription Kit (Applied Biosystems), and the resulting cDNA was analyzed by quantitative PCR using PowerTrack SYBR Green Master Mix (Applied Biosystems) on a QuantStudio 5 Real-Time PCR System (Applied Biosystems). Relative gene expression was calculated using the 2^−ΔΔCt^ method and normalized to the housekeeping gene *Hprt*. Primer sequences are listed in table S2.

### CART19 mouse model

#### Retrovirus production

One MSGV (Murine stem cell virus-based splice-gag vector) encoding mouse CD19-directed 4-1BB co-stimulated CD3ζ CAR construct (muCAR19-MSGV) was used. Murine gamma retrovirus from MSGV vectors were generated from Platinum-E (Plat-E, Cell Biolabs) cells. Plat-E cells were cultured in DMEM supplemented with 10% FBS, 1% GlutaMAX, 1% HEPES, and 1% penicillin/streptomycin (DMEM10) + 1 μg/mL puromycin + 10 μg/mL blasticidin. 24 h prior to transfection, 8 ×10^6^ Plat-E cells were seeded per T175 culture flask, and the medium was exchanged with DMEM10 (without puromycin or blasticidin). On the day of Plat-E transfection (∼70-80% confluency), a mixture of Lipofectamine 2000 (60 μL, Invitrogen) and muCAR19-MSGV retroviral construct (15 μg) was diluted in 3 mL Opti-MEM Reduced Serum medium (Thermo-Fisher) per retroviral production flask and transferred to said flasks. At both 48-and 72-h post-transfection, media containing virus was collected, and filtered (0.45 μm). The viral supernatant from 48 and 72 h were combined, flash-frozen, and stored in-80°C.

#### Mouse CAR T cell manufacturing

Mouse CAR T cells were generated using a 5 to 6-day protocol. T cells were isolated from CD45.1 BALB/c mice using the Pan T Cell Isolation Kit II (Miltenyi Biotec) and activated with anti-mouse CD3/CD28 Dynabeads (Invitrogen) at a 2:1 bead-to-cell ratio. In parallel, 24-well plates were coated with RetroNectin (Takara Bio, 1:50 in PBS) overnight at 4°C. The following day, wells were blocked with 2% BSA for 30 min, after which muCAR19-MSGV retrovirus was added, centrifuged at 2,000 × g for 1 h at 4°C, and incubated at 37°C for 30 min. Activated T cells were then transferred to the retrovirus-coated wells (0.5–1 × 10⁶ cells per well). Cells were cultured for 5 to 6 days in RPMI supplemented with 10% FBS, 1% GlutaMAX, 1% HEPES, and 1% penicillin/streptomycin (R10), with 0.05 mM 2-mercaptoethanol, 10 ng/mL mouse IL-2, and 10 ng/mL mouse IL-7, with fresh cytokines added daily. On day 5 or 6, Dynabeads were removed, and CAR expression was assessed by flow cytometry using an anti-G4S stain (Cell Signaling Technology). Expanded mouse CAR T cells were resuspended in 1x PBS for infusion into mice.

#### Syngeneic Mouse Models

To establish the A20 mouse lymphoma subcutaneous model, 2 × 10^6^ A20 cells were prepared in a 100 μL solution (50% PBS, 50% Matrigel, Corning) and implanted under the skin of the right flank in 8-to 10-week-old BALB/c mice. Eighteen days post-implantation, one day prior to CAR T cell transfer, mice were administered 100 mg/kg cyclophosphamide via intraperitoneal injection. On day 19, mice were randomized (subcutaneous tumors had reached 30–300 mm^3^) and infused with 2 ×10^6^ CAR^+^ T or an equal number of untransduced control T cells through tail vein. Tumor growth and mouse body weight were monitored twice per week. Animals were monitored for signs of disease progression and distress affecting gait or normal posture, as evidenced by >20% loss in body weight, excessive fur loss, diarrhea, conjunctivitis, disease-induced hind limb paralysis. Mice were euthanized when sign of overt disease was observed or tumor volume reached 2,000 mm^3^ or one diameter is larger than 2 cm.

#### Flow cytometry of splenic and tumor-infiltrating CAR T cells

Spleens and tumors were harvested 6 days after CART19 cell infusion. Spleens were passed through 70-μm cell strainers, followed by red blood cell lysis using ACK lysis buffer (Gibco). Cells were resuspended in 4 mL of FACS buffer, and 100 μL of the splenocyte suspension was used for staining.

A20 tumors were carefully dissected and weighed. Approximately 300 mg of tumor tissue was used for isolation of tumor-infiltrating lymphocytes (TILs). Tumors were minced and digested in 2 mL of digestion medium containing DMEM, 10% FBS, 50 U/mL penicillin and 50 mg/mL streptomycin, 20 mM HEPES, 0.5 mg/mL collagenase D, and 0.01 mg/mL DNase for 30 min at 37°C with agitation. Collagenase was inactivated by adding 40 μL of 0.5 M EDTA. Digested tissues were triturated using a P1000 pipette and passed through a 70-μm cell strainer. TILs were enriched by density-gradient centrifugation using 40% and 80% Percoll (Sigma-Aldrich). Cells were collected from the 40%–80% Percoll interface after centrifugation at 900 × g for 22 min with no brake.

For intracellular cytokine analysis, cells were stimulated with Cell Stimulation Cocktail plus protein transport inhibitors (eBioscience) for 3.5 h at 37°C. Cells were stained with LIVE/DEAD Blue Fixable Viability Dye (Invitrogen), followed by Fc receptor blockade using TruStain FcX PLUS (BioLegend). Surface staining was performed using antibodies against CD45, CD3ε, CD4, CD8α, CD45.1, CD45.2, and the CAR G4S linker (table S3). Cells were then fixed in Fixation Buffer (BioLegend) for 20 min at room temperature and stained with antibodies against IFN-γ, TNF-α, granzyme B, and IL-2 (table S3) using Intracellular Staining Perm Wash Buffer (BioLegend; Cat. No. 421002). Samples were acquired on an Aurora flow cytometer (Cytek Biosciences) and analyzed using FlowJo (version 10.10.0). Gating strategies are shown in fig. S12D.

### Serum cytokine analysis

Blood was collected by retro-orbital bleeding at baseline and on days 1, 4, and 7 after CART19 infusion. Serum concentrations of IFN-γ and TNF-α were quantified using the LEGENDplex Mouse Inflammation Panel (13-plex; BioLegend), as described above.

#### 16S rRNA sequencing and data analysis

Fecal samples were collected and stored at −80°C until DNA extraction. Genomic DNA was extracted using the DNeasy PowerSoil Pro Kit (Qiagen) according to the manufacturer’s instructions. DNA samples were submitted to SeqCenter, LLC. (Pittsburgh, PA, USA) for library preparation and sequencing. Libraries were prepared using the Quick-16S NGS Library Prep Kit (Zymo Research) with phased primers targeting the V3/V4 regions of the 16S gene. Following library cleanup and normalization, samples were sequenced on a NextSeq 2000 using a 600-cycle flow cell to generate 2 × 301-bp paired-end reads. Base calling, demultiplexing, quality control, and adapter trimming were performed using bcl-convert (v4.2.4). Primer-dimer sequences and other PCR artifacts were filtered from the resulting FASTQ files using the following criteria: read length > 150bp, PolyN strings < 10 sequential Ns; PolyG strings < 150 sequential Gs.

Taxonomic classification, diversity analyses, and stacked bar plot generation were performed using QIIME 2 (v2024.10) (*48*). Beta diversity was assessed using weighted UniFrac distances calculated from amplicon sequence variant (ASV)-level abundance tables. Permutational multivariate analysis of variance (PERMANOVA) was performed using the adonis2 function in the vegan R package (v2.6-4) (*49*). The model included time point, infection status, and cage as variables. Marginal effects were calculated using the by = “margin” option. Differential abundance analysis was performed using the DESeq2 (v1.42.1) (*50*). Differentially abundant taxa between day −7 and day 28 were identified separately for each cage. Taxa with an adjusted *P* value < 0.05 were considered significantly altered.

### Blood bulk RNA sequencing and data analysis

Blood was collected at 4 weeks after IMPACT4 exposure by cardiac puncture. RNA was extracted using the QIAamp RNA Blood Mini Kit (Qiagen), followed by on-column DNase treatment according to the manufacturer’s instructions. RNA samples were submitted to SeqCenter, LLC (Pittsburgh, PA, USA) for library preparation and sequencing. Libraries were prepared using the Illumina Stranded mRNA Prep kit and sequenced on a NovaSeq X Plus platform to generate 150-bp paired-end reads. Demultiplexing, quality control, and adapter trimming were performed using bcl-convert (v4.2.4). Reads were aligned using STAR (v2.7.10b) (*51*), and gene-level expression was quantified using RSEM (v1.3.3) (*52*).

Differential expression analysis was performed using DESeq2 (v1.42.1) (*50*) after removal of hemoglobin and mitochondrial genes. For pathway analysis, genes were ranked by log2 fold change, and gene set enrichment analysis (GSEA) was performed using the GSEA function in clusterProfiler (v4.10.1) (*53*). Gene Ontology Biological Process gene sets were obtained from the Molecular Signatures Database using msigdbr (v26.1.0) (*54*). Gene sets with adjusted P values below 0.05 were considered significantly enriched. For visualization, significantly enriched immune-related terms were selected and grouped into functional categories.

For cross-species transcriptional comparisons, the top 400 upregulated genes in either IMPACT4 or SPF mice were selected as transcriptional signatures. Mouse genes were mapped to human orthologs using babelgene (v22.9) (*55*), retaining the top-supported ortholog mapping supported by at least three databases. Human adult and cord blood PBMC microarray data were obtained from GSE27272 (*56*) and analyzed using limma (v3.58.1) (*57*). Human genes were ranked by moderated t-statistics generated using empirical Bayes modeling. Enrichment of the mapped mouse transcriptional signatures within the ranked human dataset was tested using fgseaMultilevel function in fgsea (v1.28.0) (*58*).

### Immunofluorescence and Immunohistochemical staining

For collection of the lungs, liver, kidneys, and uterus, mice were euthanized and perfused with cold PBS followed by 4% paraformaldehyde. The intestine was dissected, flushed with cold PBS, opened longitudinally, and rolled from the distal end to generate Swiss rolls. For skin collection, the dorsal region was shaved, and a square section of dorsal skin was dissected and pinned flat on a dissection tray for fixation. Tissues were fixed in 4% paraformaldehyde and transferred to 70% ethanol before paraffin embedding and sectioning. Formalin-fixed, paraffin-embedded tissue sections were used for immunofluorescence (IF) and immunohistochemical (IHC) staining.

Multiplex IF and IHC staining were performed using a Leica BOND RXm automated platform. For both assays, sections were dewaxed, rehydrated, subjected to antigen retrieval in BOND Epitope Retrieval Solution 2 (Leica, AR9640), and treated with 3% H₂O₂ for 10 min at room temperature to quench endogenous peroxidase activity. TBS (Leica, AR9590) was used for washing between steps. Negative controls were generated by replacing the primary antibodies with irrelevant isotype-matched rabbit antibodies.

Multiplex IF staining was performed as previously described (*59*) using the Opal Automation Multiplex IHC Detection Kit (Akoya Biosciences, NEL830001KT) with a Leica BOND Research Detection System (DS9455). Antigen retrieval was performed for 20 min at 98°C. Sections were blocked with Opal Antibody Diluent/Block solution (Akoya Biosciences, ARD1001EA) for 30 min at room temperature, which was also used to dilute the primary antibodies. Antibodies against CD4 (Abcam, ab183685; 1:1,500) and CD8α (Cell Signaling Technology, 98941; 1:1,000) were applied for 45 min at room temperature, followed by a biotin-free polymeric detection system containing horseradish peroxidase (HRP)-conjugated anti-rabbit antibody for 25 min. Signals were developed using the TSA reagents Opal 520 (Akoya Biosciences, FP1487001KT) and Opal 570 (Akoya Biosciences, FP1488001KT) for 10 min. Slides were counterstained with Spectral DAPI (Akoya Biosciences, FP1490) and mounted with Fluoromount-G (SouthernBiotech, 100-01).

IHC staining was performed using the BOND Polymer Refine Detection Kit (Leica, DS9800). Antigen retrieval was performed for 20 min at 95°C, followed by blocking with PowerVision IHC/ISH Super Blocking Solution (Leica, PV6122) for 30 min at room temperature. Antibodies against β-catenin (Cell Signaling Technology, 9582; 1:100) and Ki-67 (Thermo Scientific, RM-9106-S; 1:400) were diluted in SignalStain Antibody Diluent (Cell Signaling Technology, 8112) and applied for 45 min at room temperature. A biotin-free polymeric detection system (Leica, DS9800) containing HRP-conjugated goat anti-rabbit IgG was applied for 25 min, and immunoreactivity was visualized using 3,3′-diaminobenzidine (DAB). Slides were counterstained with hematoxylin, dehydrated through a graded ethanol series, cleared in xylene, and permanently coverslipped with a resinous mounting medium using a ClearVue coverslipper (Thermo Scientific).

Stained slides were scanned using a Leica Aperio Versa 8 high-resolution automated slide scanner. Images were processed and analyzed using ImageScope (version 12.4.6) and ImageJ2 (*60*) (version 2.16.0).

### Quantification and statistical analysis

Statistical analyses were performed using GraphPad Prism (version 11.0.0), except for sequencing data, which were analyzed using R (version 4.3.3). Statistical details, including the exact sample size (n), statistical test used, and significance levels, are provided in the corresponding figures and figure legends. Statistical tests were selected based on the distribution and variance characteristics of the data. The value of n represents the number of biologically independent samples analyzed. Statistical significance is denoted as follows: \**P* < 0.05; \*\**P* < 0.01; \*\*\**P* < 0.001; \*\*\*\**P* < 0.0001; ns, not significant.

